# Time-varying synergy/redundancy dominance in the human cerebral cortex

**DOI:** 10.1101/2024.06.14.599102

**Authors:** Maria Pope, Thomas F. Varley, Olaf Sporns

## Abstract

Recent work has emphasized the ubiquity of higher-order interactions in brain function. These interactions can be characterized as being either redundancy or synergy-dominated by the heuristic O-information [1]. Though the O-information can be decomposed into local values to measure the synergy-redundancy dominance at each point in a time series [2] no such analysis of fMRI dynamics has been carried out. Here we analyze the moment-to-moment synergy and redundancy dominance of the fMRI BOLD signal during rest for 95 unrelated subjects. We present results from several interaction sizes. The whole brain is rarely synergy-dominated, with some subjects never experiencing a whole-brain synergistic moment. Randomly sampled subsets of many sizes reveal that subsets that are the most redundancy dominated on average exhibit both the most synergistic and most redundant time points. Exhaustive calculation of the optimally synergistic and optimally redundant triads further emphasizes this finding, with recurrent nodes frequently belonging to a single coherent functional system. We find that when a triad is momentarily synergistic, it is often split between two instantaneously co-fluctuating communities, but is collectively co-fluctuating when it is momentarily redundant. After optimizing for synergy and redundancy in subsets of size five to seventy-five, we show that this effect is consistent across interaction sizes. Additionally, we find notable temporal structure in all optimized redundant and synergistic subsets: higher order redundant and synergistic interactions change smoothly in time and recur more than expected by chance.

## I. INTRODUCTION

There has been substantial recent interest in higher-order interactions in brain activity [3–8], and in particular, in distinguishing between redundant and synergistic higher order interactions using information theory [9–16]. Redundant higher order interactions are those interactions between three or more brain regions in which information is copied across regions. Higher order synergistic interactions are interactions between three or more brain regions in which there is some information at the level of the whole that is not reducible to the activity of individual brain regions. Several studies have revealed that traditional pairwise methods of studying interactions between regions, such as functional connectivity analyses, almost exclusively capture redundancy and are blind to synergistic interactions [10, 12, 17]. However, synergistic interactions are thought to be highly relevant to information modification [18] and computation in complex systems[19, 20] and have begun to be fruitfully identified in neural systems. In neuronal recordings and simulations, statistical synergy has been used to characterize distributed computation [20–25], and the balance of synergy and redundancy predicts success during an auditory discrimination task [16]. In recordings of neuronal dynamics in macaques engaged in complex behaviors, the distributions of synergies and redundancies through-out the population changes in response to the varying demands of different tasks [26]. In humans, synergistic interactions are clinically relevant and implicated in Alzheimer’s disease [27], stroke recovery [28], schizophrenia [29], and Autism Spectrum Disorder [30]. They change characteristically during healthy aging [14, 15], and are clinically manipulable through transcranial ultrasound stimulation [31]. Additionally, several studies have proposed that synergistic interactions play a key role in human cognition and consciousness [11, 32–34].

The O-information, a heuristic measure of redundancy/synergy dominance[1, 35], has been instrumental in revealing the presence of higher order interactions in the cortex as measured with blood oxygen-level dependent functional magnetic resonance imaging (BOLD fMRI) [10, 17, 36], in particular because it scales to handle interactions of many components, unlike other statistical approaches, such as the partial information decomposition [9]. Like many other information theoretic metrics, the O-information can be written as an expected value over all of a system’s realized states. This expected value can be unrolled into a local value corresponding to each particular state of the system [19]. Because the system enters a new state at every point in time, these local values can be used to measure the synergy/redundancy dominance of the instantaneous state of the system. This approach to localized information theory was first pioneered by Lizier[19, 37], and has been used to reveal dynamic information structures in many systems, such as cellular automata [38, 39], Boolean networks [40], neuronal populations [41], schools of fish [42], and music [2].

We note that this approach refers to temporal localization, and is not to be confused with spatial localization, which examines lower-order contributions to the higher-order O-information [43], a technique which has already been fruitfully applied to BOLD fMRI data [36]. The time-localized O-information, on the other hand, can be used to conduct an analysis of the time-varying structure of higher order redundant and synergistic interactions, and has not yet been applied to BOLD fMRI data.

Quantifying time-varying interactions has been of long-standing interest in network neuroscience [44], and developing tools to understand how brain activity varies in time is key to relating it to cognition and behavior, both temporal phenomena. Recently the interest in time-resolved interactions has intensified with the use of instantaneous co-activation metrics[45]. Measures of co-activation describe how instantaneous pairwise interactions change through time and can be used to increase subject-specific identifiability[46], are known to align with salient time points during movie-watching[47], and change characteristically during healthy aging[48]. Using the local O-information, we are able to extend this field of research to study even larger interactions, and to specifically characterize their information content. Despite co-existing interests in synergy and time-varying connectivity, most studies of synergistic interactions to date have left unexamined how these interactions evolve through time. While the approach of Luppi et al. [32] incorporates a time-resolved decomposition of the integrated information, it was focused on information shared between individual nodes’ past and future, rather than interactions between several nodes. A full analysis of time-varying synergy and redundancy dominance that respects the irreducible nature of synergy has yet to be performed. The local O-information is a good candidate measure for a first analysis of time-varying synergy and redundancy both because it is scalable to large interaction sizes, and because the efficacy of its expected value on BOLD fMRI data has already been demonstrated. While some work has already been done validating the local O-information in Bach choral music in ref. [2], and in applications to physiological signals and EEG data[49], ours is the first study to apply it to BOLD fMRI data. We hope to provide a thorough analysis of ongoing higher-order dynamics during resting state fMRI across several sizes of interaction and with a particular eye to characterizing the type of states that result in a negative (synergistic) local O-information.

## II. METHODS

### A. Dataset and Preprocessing: HCP

We studied the time-varying synergy/redundancy dominance of functional magnetic resonance imaging (fMRI) data taken from 100 unrelated subjects of the Human Connectome Project [50]. fMRI measures the ongoing blood oxygenation level in the cerebral cortex, which increases with neuronal activity [cite]. Increases in blood oxygenation happen on a slower time scale than the neuronal activity itself, following a haemodynamic response function that results in a smooth time series well-approximated by a Gaussian distribution. The dataset consists of four scans per participant, each with a duration of 14:33 minutes, performed across two separate days. Informed consent was provided by all participants and data collection protocols were approved by the Institutional Review Board at Washington University. All imaging scans were performed at rest; the participants were instructed to look at a fixation cross and allow their minds to wander. The data was minimally preprocessed according to [51] at the time of download.

The scans were collected with a Siemens 3T Connectom Skyra with a 32-channel head coil. A gradient-echo echo-planar imaging (EPI) sequence was used with the following parameters: TR = 720ms, TE = 33.1ms, flip angle = 52°, isotropic voxel resolution = 2mm, and multiband factor = 8.

Participants were included in the current study based on several data quality factors established prior to the study. Four subjects were excluded due to too much motion in the scnner. They exceeded 1.5 times the interquartile range of the distribution of the mean and mean absolute deviation of the relative root mean square (RMS) for motion calculated across all four scans. Another subject was excluded because of software errors during processing of their diffusion MRI. The subjects included were 56% female and 22-36 years old, with a means age of 29.29 *±* 3.66.

Upon download the data was in the form of an ICA+FIX time series in the CIFTI grayordinate coorodinate system, and we performed further preprocessing steps. The data was global signal regressed and band pass filtered (0.008 Hz to 0.08 Hz [52]). Confound regression and filtering were also performed, and then the first and last 50 TRs of the data were discarded. This resulted in time series with a final length of 13.2 minutes captured by 1100 TRs, or time points. To parcellate the data into brain regions, the surface-level data was then averaged at each time point within each parcel of the 200-parcel Schaefer atlas in CIFTI coordinates [53], resulting in a time series of 200 brain regions (nodes) per subject.

For all of the analyses in this study, all individual subject data was concatenated into a single master subject. This was done to take full advantage of the entire dataset when estimating the large multivariate joint distributions required for the O-information, and to minimize as much as possible the noise resulting from undersampling a large distribution. Since each subject contributed four scans each with 1100 time points, this resulted in a final time series of 200 nodes and 418,000 time points. This time series was z-scored so that each node had a mean activity level of zero.

### B. Dataset and Preprocessing: MICA

Results were replicated in a second dataset from 50 unrelated participants, undergoing one seven minute scan each. As in the HCP dataset, participants were instructed to rest while looking at a fixation cross for seven minutes. All participants provided written informed consent to data collection protocols approved by the Ethics Committee of the Montreal Neurological Institute and Hospital. Participants were scanned with a 64-channel head coil 3T Siemens Magnetom Prisma-Fit. An EPI sequence was used with the following parameters: TR = 600ms, TE = 48ms, flip angle = 52°, isotorpic voxel resolution = 3mm, and multiband factor = 6.

Preprocessing of the MICA dataset followed the protocols in reference [54]. The Micapipe data processing pipeline [55] was used for motion and distortion correction in addition to FSL’s ICA FIX tool trained with an in-house classifier. Time series were then projected onto FreeSurfer surfaces for each subject where nodes were defined according to the Schaefer parcellation [53], as in the HCP data above. This resulted in a time series with 200 nodes and 695 time points for each subject. In addition to all of the steps described in ref. [55], the data was also global signal regressed.

Again, as in the HCP data, all individual subject data was concatenated to form a single master time series of 200 nodes and 34,750 time points. This time series was also z-scored, resulting in a mean of zero for each node. All multivariate joint distributions were estimated from this time series.

### C. The O-information

In order to study the time-varying synergy and redundancy dominance in fMRI BOLD data, we used the O-information. The O-information is a heuristic measure of synergy or redundancy dominance. First introduced as the enigmatic information[35], it was later re-analyzed and clarified by Rosas and Mediano [1] who renamed it the O-information, for “organization information”. Formally, the O-information of a system **X**, Ω(**X**) is defined as:

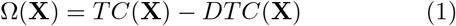

The O-information is composed of two multivariate generalizations of the mutual information, the total correlation (*TC*(**X**)) and the dual total correlation (*DTC*(**X**)). The total correlation is a measure of a system’s integration as a whole, or its collective deviation from independence. It is defined as:

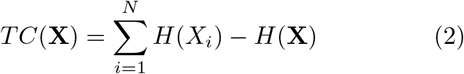

where *H*(*Xi*) is the Shannon entropy of every individual variable, and *H*(**X**) is the joint entropy of the system. The total correlation is maximal when a system is fully synchronized and is zero when all variables of the system are completely independent. The dual total correlation measures the extent to which the integration of a system is contained in the whole and not in the parts. It is given by:

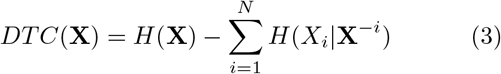

where the notation **X**^*−i*^ indicates the joint state of the system without the element *Xi*. The DTC is zero when the system is entirely synchronized and also zero when it is entirely independent. It peaks in a regime where some information is shared without being globally redundant. When the DTC is larger than the TC, the O-information is negative and we say that the system is synergy dominated. However, when the TC is larger than the DTC, then the O-information is positive and the system is redundancy dominated [1].

### D. Localizing the O-information

The approach to temporally localizing information theoretic measures was pioneered by Lizier [19], and leverages the fact that almost all information theoretic measures can be written as expected values averaged over all possible states of the system. For instance, the entropy in the measures above can be written as *H*(*X*) = E[*−log*2*p*(*x*)]. Removing the expected value leaves the “surprisal,” or local entropy of a specific state *x* of the variable *X*. The surprisal can be used as a temporal localization of the entropy since at each point in time, a variable takes on a specific state, and so has the surprisal value associated with that state. Intuitively, this is a measure of how surprised an observer is to see that specific state, given the probability distribution of states for the variable. This is written as:

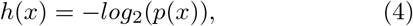

where *x* is a specific state of the variable *X*, and *p*(*x*) is the probability that *X* adopts that state. (Here we use the lower-case to denote local measures.)

The joint entropy can be similarly localized to *h*(***x***) = *−log*2(*P* (***x***)). Total correlation, dual total correlation, and O-information can all be written in terms of joint and conditional entropies, so we can write the localized version of each metric in a similar way. The total correlation becomes:

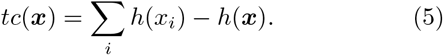

The dual total correlation is written as:

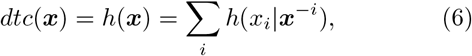

and the O-information is a combination of these two measures:

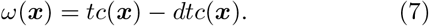

The local O-information is the measure explored primarily in this paper and is used to obtain a time-point by time-point measure of synergy/redundancy dominance. It is important to note that the local forms of the total correlation and dual total correlation do not have all the same properties as their expected value counterparts. Most importantly, they can take on negative values. This means that the local O-information may become negative in two ways: either the local total correlation is strongly negative, or the local dual total correlation is strongly positive. We do not believe that this changes the interpretation that a negative value indicates synergy dominance. However, this is discussed in more detail in Supplemental Text 1.

### E. Gaussian Information Theory

Since it has been demonstrated that fMRI BOLD data is well-approximated by multivariate Gaussian distributions [56, 57], all information theoretic calculations in this paper were made using closed-form estimators of Gaussian entropy and joint entropy. The entropy of a single Gaussian random variable *Xi* with standard deviation *σ* is:

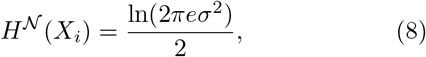

measured in nats. The joint entropy of a multivariate Gaussian random variable, where **X** = *X*1, *X*2, …*XN*, is:

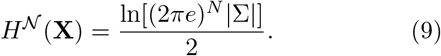

|Σ| is the determinant of the covariance matrix and *N* is the number of dimensions.

Both of these can be localized by taking the natural log of the probability density of a given state of the variable, such as:

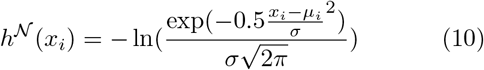

for the local entropy or surprisal where *µ* and *σ* are the mean and standard deviation of *xi* respectively, and

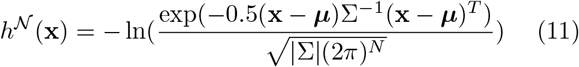

for the local joint entropy, where **x** is the joint state *x*1, *x*2, …*xN*, ***µ*** is a vector of the mean of variables *x*1, *x*2, …*xN*, *N* is the dimensionality of the system, Σ is the covariance matrix, and |Σ| is the determinant of the covariance matrix. Both local forms are also measured in nats.

This estimator was used to calculate all information theoretic measures introduced in the previous section.

### F. Sampling Methods

The space of possible higher order interactions in a 200-dimensional system is massive. It is impossible to thoroughly investigate all interactions of all sizes, since we have no *a priori* assumptions about the most important orders of interaction. The irreducibility of synergistic interactions also means that each size of interaction may be unique from other sizes, and so findings at any size cannot necessarily be extrapolated to other sizes. To begin to overcome this problem, we sampled subsets of nodes using four different strategies.

First, we treated the whole brain as a single large interaction and calculated a single local O-information time series for all time points. Second, we randomly sampled subsets of size three to 25. 10,000 subsets were sampled for each size, and a local O-information time series was calculated for each subset. Primarily these were used to assess how the local O-information values related to the same subset’s O-information expected value. Third, we found the maximally synergistic and maximally redundant triad independently for each time point. This was done by exhaustively calculating the local O-information time series for every possible triad (1,313,400 triads, 418,000 time points per triad) and retaining the triad with the minimum (for synergistic) or maximum (for redundant) local O-information at each time point (see schematic in Figure 2a). This process produced a time series of subsets as well as a time series of the optimal local O-information at each time point. We repeated the analysis to find the most synergistic tetrad at every time point (out of 64,684,950 possible tetrads). Redundant tetrads could not be included because of an error in saving the data. Optimal triads were analyzed for nodal autocorrelation and recurrence, methods which are outlined below. Finally, for larger subsets, calculation of all possible subsets is prohibitive, so we performed an optimization using a simulated annealing algorithm [10, 58]. The algorithm was run independently on every time point and for subset sizes five to seventy-five. This process is described in more depth below.

**FIG. 1.**
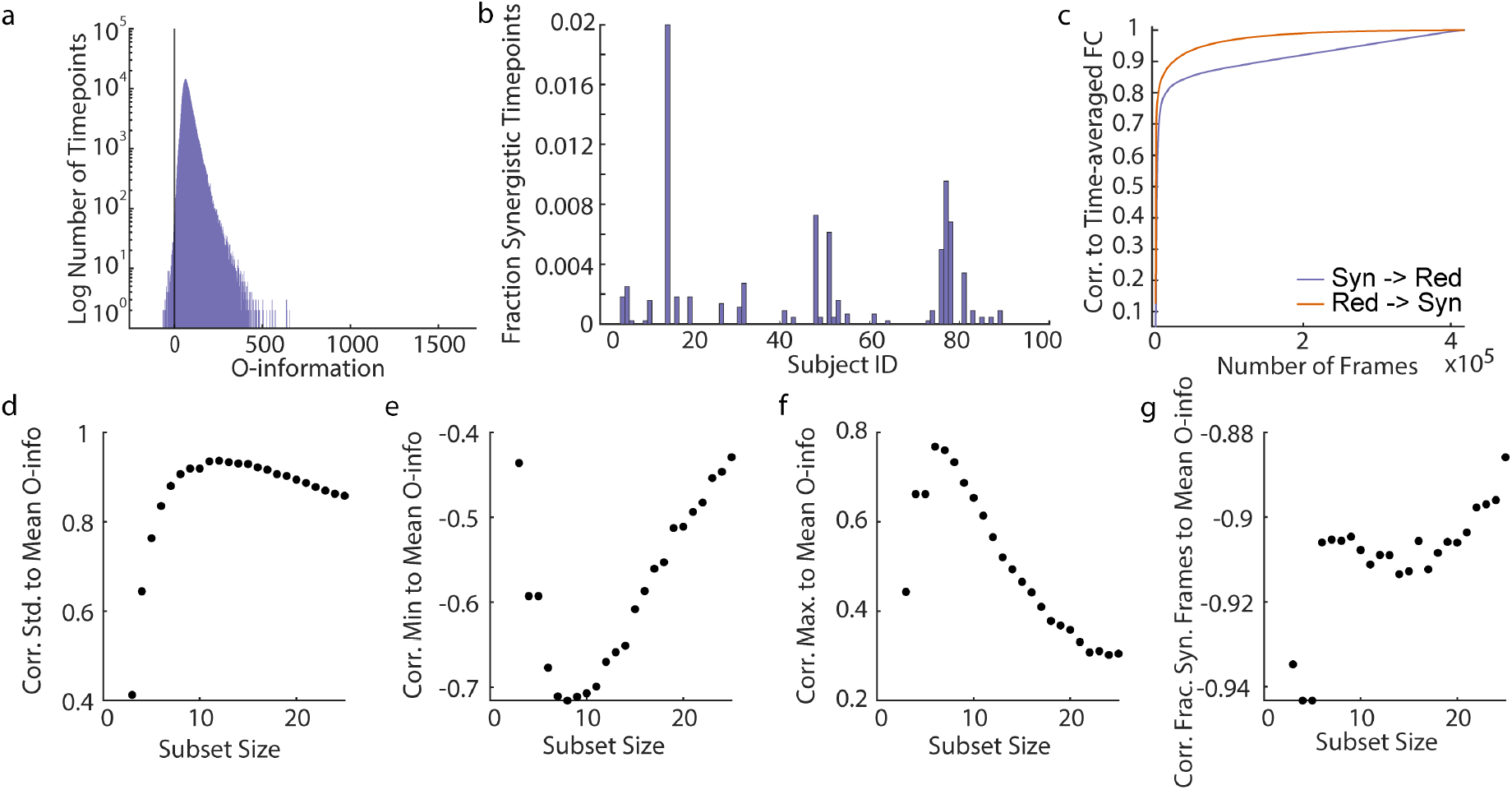
The whole brain is rarely synergy dominated. Redundancy-dominated subsets have the most synergistic and the most redundant moments. **a**. A histogram of all local O-information values for all 200 brain regions, calculated on all subjects and all scans. A grey line marks zero. **b**. The number of whole-brain synergy dominated time points by subject. Many subjects do not ever experience a whole-brain synergistic time point and it is a rare event for those that do. **c**. The correlation to time-averaged functional connectivity of the sequential averaging of BOLD edge time series frames ordered by their local O-information value. Time-averaged functional connectivity is much more quickly approximated by averaging positive (redundancy dominated) local O-information time points first. **d.-g**. The correlation of local O-information summary statistics (**d**. standard deviation, **e**. minimum value, **f**. maximum value, **g**. fraction of synergistic time points) to the expected value of the O-information of the subset for 10,000 randomly sampled subsets of each subset size. The strong correlations in **d.-f**. indicate that randomly sampled, redundancy-dominated subsets tend to have a larger dynamic range than randomly sampled synergy dominated subsets.

**FIG. 2.**
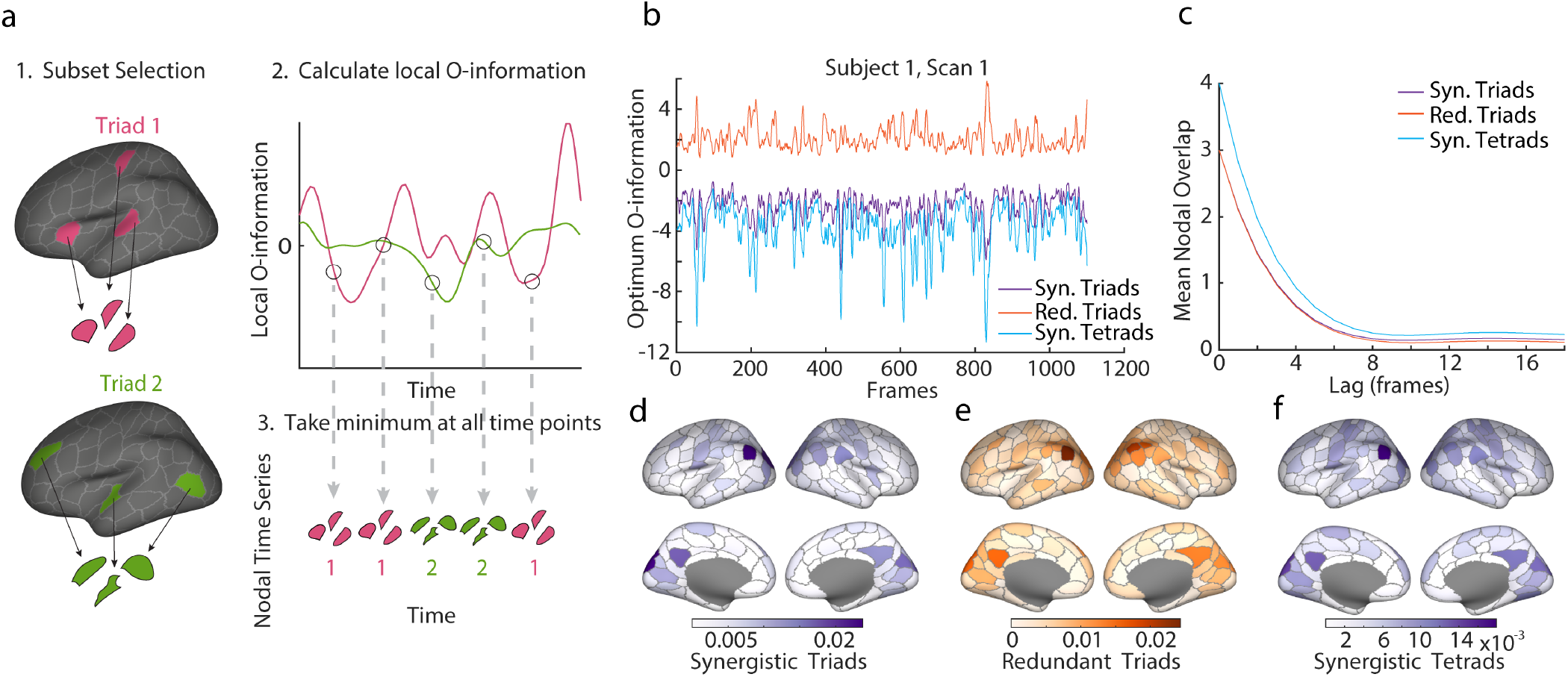
Calculation of maximally synergistic triads and tetrads and maximally redundant triads. **a**. A schematic illustrating triad and tetrad selection. The local O-information was calculated for all triads/tetrads and then compared independently at each time point. The identity of the minimal/maximal triad/tetrad and the local O-information value of that subset was retained for each time point. **b**. A sample time series of the maximum and minimum local O-information found at each time point, which may act as upper and lower bounds of the local O-information. **c**. The average number of nodes shared between time points at varying frame lags. This serves as an autocorrelation for the nodal time series. **d.-f**. The frequency of participation in synergistic triads and tetrads and redundant triads plotted on the surface of the cortex. A wide variety of brain regions participate, but not uniformly.

### G. Analysis of Optimal Triads

Once maximally synergistic triads were identified at every time point, we performed several analyses to assess the temporal structure of synergistic and redundant interactions. First, to understand if interactions changed smoothly in time or jumped around the cortex, the autocorrelation of the triad identity was measured. We counted the number of overlapping nodes between every pair of time points at successively longer lags. This value was then averaged over all time point pairs to produce a single autocorrelation value per time lag. Second to see if synergistic and redunant interactions were repeated across time, a recurrence matrix was created by counting the number of nodes in common between all time points within an individual subject. Recurrence matrices were calculated individually for each subject, resulting in 95 matrices each 4400 x 4400 time points. To better understand the structure of recurrent time points and to identify recurrent states, multi-resolution consensus clustering [59] was applied separately to each of these matrices. Multi-resolution consensus clustering is an algorithm that searches for community partitions across modularity resolution levels, creating a single, data driven partition that leverages the agreement between many partitions. In this algorithm, modularity maximization was first performed on a series of 100 resolution parameters producing partitions containing between 2 and N modules. A second, more fine-grained stepping of the resolution parameter through 1000 values yielded a set of partitions which were then aggregated into a node-by-node coclassification matrix with values between 0 and 1. After subtraction the analytic null detailed in [59], we applied consensus clustering to the resulting matrix with *τ* = 0. This process was repeated for every subject’s recurrence matrix.

We used the clusters of nodes identified within individual subjects to create group states. To create group clusters, we first identified the nodes involved in each subject-specific state. The node membership for each subject-level cluster was binarized to make a nodal-participation vector that stored whether or not a node was ever present in the cluster. (While this method loses some information about the frequency of nodal participation in a subject-level cluster, we argue that there was so little range in these values to begin with (see Supplementary Figure 2) that it does not affect the main results and makes computation substantially easier.) We created a similarity matrix between subject states by taking the Jaccard similarity 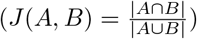 between every subject-specific nodal participation vector. We applied multi-resolution consensus clustering to the resulting matrix using the same process described above. For both synergistic and redundant triads, this produced three large group-aggregated states, and a unique assignment of each time point to a single group-level state. The time series of group-level states was used to create state transition matrices by counting the frequency of transition from one group-level state to another within every scan.

Finally, triads and tetrads were analyzed according to their instantaneous node-level activity. For this analysis only, nodal activity was thresholded at zero to create two communities of instantaneously co-fluctuating nodes, one with positive z-scores and the other with negative. We refer to this as the bipartition [60], and it has been demonstrated that this binarization preserves much of the information available in the original time series [60]. We then count how many nodes fall on the same side of the bipartition for every triad and tetrad.

### H. Optimization and Analysis of Larger Subsets

Because calculation of all possible larger subsets is prohibitive, we optimized subsets for positive and negative local O-information. To do this, we used simulated annealing [58], a probabilistic optimization algorithm that was used to optimize larger subsets for positive and negative local O-information. Simulated annealing was run independently for every subset size on every time step, with subset sizes of 5-75, counting by 5. A total of 15 (subset sizes) *×* 418,000 (time points) *×* 2 (redundancy/synergy) simulated annealing operations were performed. On each trial, the simulated annealing algorithm was seeded with the optimally synergistic tetrad, or optimally redundant triad, corresponding to that particular time point, and additional nodes were chosen at random to bring the initial subset to the desired subset size. Simulated annealing then proceeds on each step to generate a new subset by randomly swapping in new nodes to the subset. The number of nodes to swap can be no more than the size of the subset and is sampled from a Gaussian distribution centered at zero (the absolute value of the nearest integer is chosen). The local O-information of the new subset is computed, and the new subset is kept if the local O-information is lower (for synergy, higher for redundancy) than the original subset, or if a random number drawn from a uniform distribution between 0 and 1 was less than the threshold exp(*−* ((*Cn − C*)*/Tc*)), where *Cn* is the local O-information of the new subset, *C* is the local O-information of the current subset and *Tc* is the current temperature. The current temperature decays to a fraction of the initial temperature at each annealing step, given by:

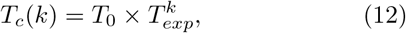

where *k* is the number of steps completed, *T*0 is the initial temperature (set to 1), and *Texp* determines the steepnees of the temperature gradient. The progressive decreasing of the temperature as the algorithm runs allows for more stochasitc exploration of the optimization landscape in the beginning, but restricts it to more deterministic local gradient descent toward the end of the run. Each run was allowed a total of 10,000 annealing steps. We note that for all multivariate spaces explored, 10,000 steps explores only a very small fraction of the possible subsets within the space. For this reason we cannot make claims about exhaustive solution optimality.

The subsets found by simulated annealing were also analyzed for autocorrelation, recurrence and node-level activity. The node-level bipartition analyses proceeded exactly as described for the triads and tetrads. However, to identify autocorrelation and significant recurrences, we created a null distribution of the overlap of pairs of randomly sampled subsets. For each subset size, 2,000,000 subsets were sampled to produce a null distribution of 1,000,000 overlap values. To be considered an autocorrelated time point or a significant reucrrence, time points had to have significantly greater overlap than this null distribution with Bonferroni-corrected p-values. The number of significant time points at each time lag was averaged to create an autocorrelation function with a single expected overlap value per time lag.

## III. RESULTS

We sought to examine time-varying synergy and redundancy dominance using the time-localized O-information (see Methods) for 95 unrelated subjects from the Human Connectome Project [50] (see Methods), using the Schaefer parcellation [53] of 200 brain regions spanning the entire cortical surface. However, for a 200 element system, the space of possible interactions between three or more elements is massive (a total of *∼*1.6 *×* 10^60^ possible interactions for a 200 element system). It is impossible to examine all of these interactions, and it is clear that any method one chooses to select interactions for study will have a large effect on the findings. We explored four ways of tackling this problem: treating all brain regions as a single interaction, randomly sampling brain regions, exhaustively calculating all 3- and 4-order interactions, and optimizing for synergistic interactions separately on each time point and for various subset sizes. What we hope emerges is an outline of the structure of time-varying synergistic and redundant interactions in the cortex across several orders of interaction.

### A. Whole-Brain Synergistic Moments

To begin, the local O-information of the whole brain was calculated on all time points for all subjects and all scans. We found that the whole brain is usually redundancy dominated and rarely experiences any moments with negative O-information. Figure 1a shows a histogram of the local O-information values for all time points in the dataset. Synergy-dominated time points represent only 364 time points out of 418,000 (0.087 % of time points). Additionally, whole brain synergy-dominated moments are not uniformly distributed over subjects, with many subjects never experiencing any whole brain synergy dominated moments (Figure 1b). To see if there were any consistent activity patterns during whole brain synergistic moments, we clustered the instantaneous co-activation patterns on all time points with negative local O-information using multi-resolution consensus clustering (see Methods). The clustering revealed only two major communities of co-activation patterns, both of which produced very noisy centroids (Supplementary Figure 1). This result, coupled with the rare occurrence of whole brain synergistic time points may indicate that the O-information, when applied to interaction orders larger than a sensible size for the system is particularly susceptible to noise (see Discussion and further Results). Nonetheless, the whole-brain local O-information can still serve to indicate which time points more closely replicate time-averaged functional connectivity. Sequentially averaging the instantaneous co-activation of every time point more quickly approximated time-averaged functional connectivity when moving from redundancy-dominated time points to synergistic ones than when moving from synergistic time points to redundant ones (Figure 1c) This is consistent with previous work suggesting that FC is driven more by higher-order redundancies than by higher-order synergies [10, 12].

### B. Randomly Sampled Subsets

As a first exploration of the interactions of subsets smaller than the whole brain, we randomly sampled 10,000 subsets of sizes 5 to 25 brain regions. To understand how the local O-information relates to the expected value of the O-information used in other studies of BOLD fMRI [10], we correlated several summary statistics of the local O-information to the expected value (mean) of the local O-information for each subset size. Significant correlations were found for all subset sizes and all measures. We found large positive correlations between the standard deviation and mean local O-information as well as the maximum value and mean local O-information for most subset sizes in addition to large negative correlations between the minimum value and mean local O-information (see Figure 1d-f). These correlations indicate that subsets that are redundancy dominated on average tend to have the greatest dynamic range (measured by the standard deviation of the local O-information) and experience both the most redundant and the most synergistic moments in the dataset (indicated by the maximum and minimum value of the local O-information, respectively). Finally, to gain insight into how much the time-averaged O-information may be driven by these extremal points, the fraction of synergistic time points was correlated to the mean O-information across subset sizes. A stronger negative correlation indicates that the time-averaged value is less driven by extreme values. All subset sizes had fairly strong correlation values, but the strength of the correlation tended to decrease with subset size, with a fairly large jump after subset size 6 (Figure 1g). As a whole, these results indicate that the expected value of the local O-information, the form of the measure most frequently used in studies of fMRI data, does not always give good insights into time-varying synergy and redundancy dominance.

### C. Maximally Synergistic and Maximally Redundant Triads

To explore where and when the most synergistic and most redundant interactions are taking place in the cortex, we calculated the time series of local O-information values for every possible triad of the 200 node parcellation. This dataset was used to determine the most synergistic and most redundant triad at every point in time and was repeated to identify the most synergistic tetrad at every time point. A schematic of the triad selection process and resulting time series of synergistic and redundant triads can be found in Figure 2a. This analysis produces both a time series of optimal local O-information values and a time series of optimal triads. A sample of the optimal local O-information time series for the first subject and first scan can be found in Figure 2b. These time series represent upper or lower bounds on the local O-information value for their particular subset size at every point in time, a value which fluctuates strongly and which future work may be able to use to identify particular time points of interest, but it will not be studied further here.

Our analysis identified a large number of unique triads, and implicated many brain regions at least once. Specifically, 8,112 unique triads were identified as the maximally synergistic triad at at least one point in time, and all brain regions were included in a maximally synergistic triad at least once. Similarly, 12,122 unique triads were identified as the maximally redundant triad at at least one point in time, and all brain regions except one were included in a maximally redundant triad at least once. Figure 2d-f shows the frequency of nodal participation in triads and tetrads. Since our focus was limited to only one triad per time point, the large number of unique triads indicates an abundance of ongoing synergistic and redundant interactions at rest. However, the number of triads selected still represents a very small fraction of the total space of possible triads, so, while ubiquitous and widespread in the cortex, higher order interactions are largely specific to particular triads across time and subjects. Additionally, the nodes participating in maximally synergistic and redundant interactions at the triad and tetrad level change smoothly. Figure 2c shows the mean nodal overlap between chosen subsets at time points offset from each other by different lag times. This autocorrelation reaches zero at about 8 time points, which accords with the slow nature of the hemodynamic response expected in a BOLD time series. This analysis also produces a time series of optimal local O-information values. The strongly fluctuating optimal local O-information as well as the frequency of nodal participation maps were replicated in a second independent dataset (MICA)[54] that was collected with different acquisition protocols (see Methods and Supplemental Figure 2). The lagged mean nodal overlap was the only strong difference between the two datasets, with the MICA dataset’s mean nodal overlap falling off completely by a lag of 3-4 time points. This cannot be explained by the native autocorrelation of the MICA dataset (which falls off at a lag of about 20 time points) and warrants further study.

Since the number of unique triads is much less than the total number of time points in the dataset, maximally synergistic and maximally redundant triads must recur substantially over the course of a scan. To investigate these recurrent structures, we first created a time point-by-time point recurrence matrix for each subject (a sample recurrence matrix for the first subject, see Supplementary Figure 3). We began at the subject level and performed multi-resolution consensus clustering on each of these matrices (a sample agreement matrix for the first subject can be found in Supplementary Figure 3). The number of clusters found in each subject varied, but with a mode of five clusters for maximally synergistic triads and eight clusters for maximally redundant triads (Figure 3 a and f). To create group averaged clusters, we then further applied multi-resolution consensus clustering to a matrix storing the similarity between subject level clusters. Distance matrices are shown in Figure 3 b and g. The agreement matrices resulting from this clustering for redundant and synergistic moments are also reported in Figure 3 c and h, respectively. This process gave three distinct clusters for both redundant and synergistic moments and a unique identification with one of these clusters for every time point. The frequency of nodal participation (as a fraction of total time points) in each of these three clusters highlight nodes that correspond to known functional systems [61] (Figure 3 e and j), specifically the Somatomotor, Visual, and Default Mode Networks. This indicates that these nodes are repeatedly selected as participating in the most redundant and synergistic triads, and further, nodes within these communities tend to frequently recur together. Highly similar recurrence patterns were found in the MICA data using the same data analysis pipeline (see Supplemental Figure 2). While the alignment of synergistic triads to known functional systems may seem unexpected based on ours and others’ previous work indicating that these systems are redundancy dominated [10, 12, 17], it is consistent with the results in Figure 1d-g that show that globally redundant subsets experience the most synergistic moments.

**FIG. 3.**
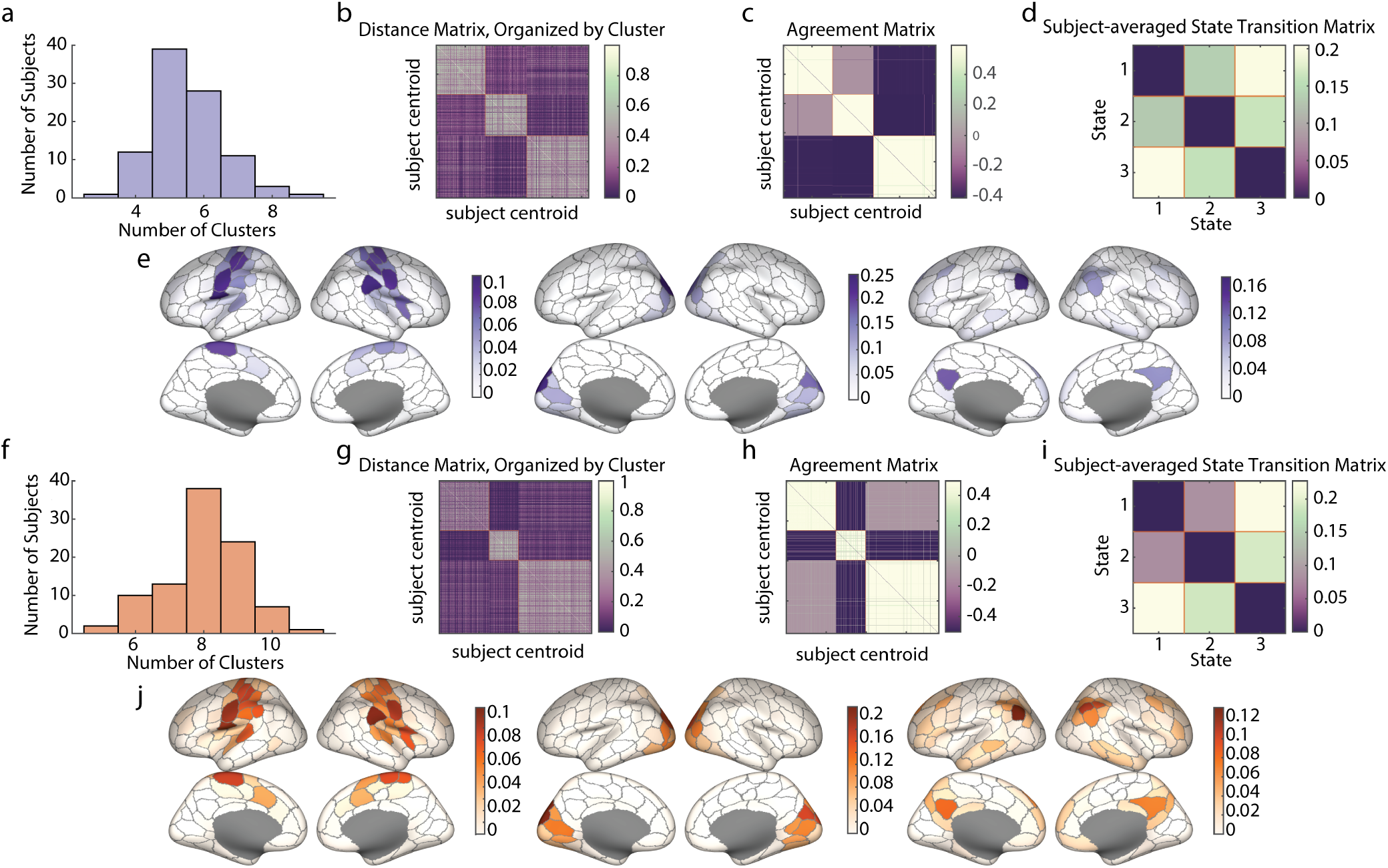
Maximally synergistic and maximally redundant triads recur. **a**.,**f**. Histograms of the number of clusters discovered in each subject during multi-resolution consensus clustering (MRCC) of triads on single subjects. Synergistic triads shown in **a**. and redundant in **f. b**.,**g**. The distance matrix between nodal participation vectors from the centroid of every subject level cluster. The matrices have been organized into group-level clusters found for synergistic triads (**b**.) and redundant triads **g. c**.,**h**. The agreement matrices for group level MRCC of the subject-level centroids, organized by resulting clusters. Synergistic triads shown in **c**. and redundant in **h**.. **d**.,**i**. State transition matrices for the group-level clusters shown for synergistic and redundant triads, respectively. **e**.,**j**. Nodal participation frequency plotted on the cortical surface for each of the three clusters found in synergisitc and redundant triads, respectively.

What makes a globally redundant subset become momentarily synergistic? To probe this further, we closely examined the relationship of the Yeo functional systems’ [61] activation to the maximally synergistic/redundant triads at each time point. First, we confirm the intuition developed in Figure 3e and j that synergistic and redundant triads indeed map onto the Yeo systems far more than is statistically expected based on system size (Figure 4a.). The maximally synergistic or redundant triad is entirely contained within a Yeo System on more than 60% of time points, which is substantially greater than expected by chance (Figure 4a). One of the key properties of Yeo functional systems is that they tend to remain integrated on long time scales, but they may not be integrated at every moment in time. We thought that their momentary distintegration may be related to synergy dominated moments, and so we examined the triads’ and tetrads’ relationship to the instantaneous bipartition of the network. The bipartition is a natural grouping of all nodes into two communities based on the sign of their instantaneous z-score. Nodes are grouped into one co-fluctuating positive community and one co-fluctuating negative community [60]. We consistently find that at the moment they are chosen, maximally synergistic triads tend to have one node that has a different sign than the other two. In contrast, when triads are interacting redundantly, they are located on the same side of the bipartition and all nodes have the same sign of their z-score (Figure 4b). This trend was also observed in the maximally synergistic tetrads (Figure 4c.). Finally, to tie the bipartition to the Yeo systems, we found that when a triad becomes synergistic and is fully contained within a Yeo system, the Yeo system itself is more likely to be split evenly across the bipartition (roughly half of its nodes have positive z-scores and half have negative), whereas the opposite tendency is true of a Yeo system that fully contains a redundant triad (Figure 4d.).

**FIG. 4.**
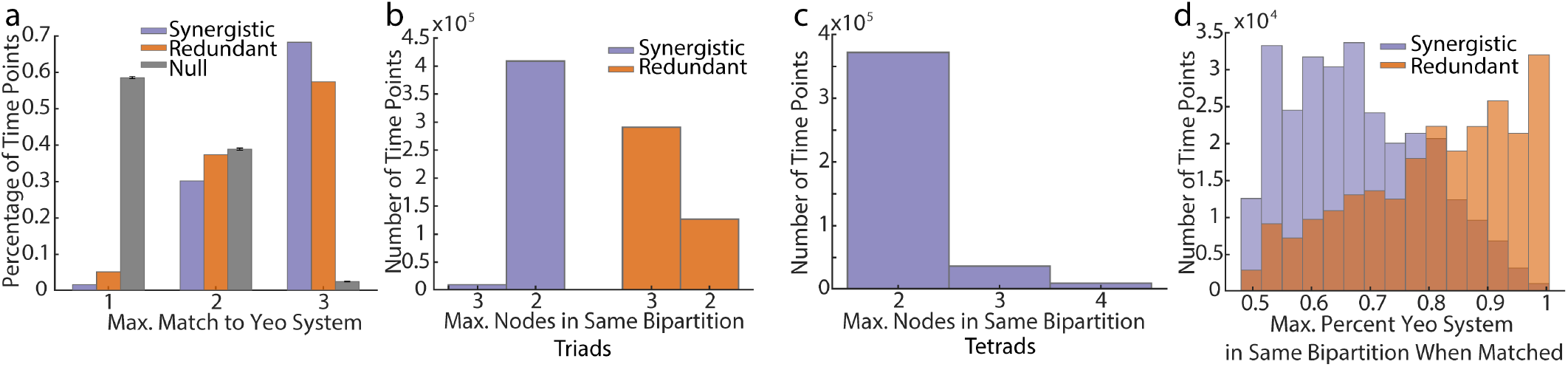
Both synergistic and redundant triads are contained within Yeo systems, but may be distinguished by their relationship to the bipartition. **a**. It is possible for a triad to match a Yeo system in three different ways: all nodes may be in different Yeo systems, two nodes may match the same Yeo system, or the triad may be entirely contained within a Yeo system. The percentage of time points corresponding to each of these matching conditions is plotted for synergistic and redundant tetrads alongside a null model built by randomly sampling triads 10,000 times. Error bars indicate the maximum and minimum values found in the null sampling. **b**. Triads can have one of two possible relationships to the bipartition (set of two nodal communities determined by the sign of their z-scores, see text). The percentage of time points corresponding to each possible bipartition condition is plotted for redundant and synergistic triads. The difference between these distributions is statistically significant. **c**. Tetrads can have three possible relationship to the bipartition (see text). The number of time points corresponding to each of these relationships is plotted for maximally synergistic tetrads. **d**. Histograms of the maximum match to either side of the bipartition of the Yeo system when the chosen triad is fully contained within that Yeo system for both redundant and synergistic triads. Histograms are statistically significantly different (Two-sample Kolmogorov-Smirnov test, KS stat = 0.4106, *p <* 1 *×* 10^*−*16^).

A rigorous mathematical explanation for the relationship between small subsets, the Yeo systems, and the local O-information was found by leveraging the combination of the particular structure of BOLD data and the conditions under which the local O information can be negative, specifically in the case of negative total correlation. We demonstrate this in Supplementary Figure 4 and Text 1. However, this explanation does not necessarily apply to larger subsets, as the ratio of total correlation magnitude to dual total correlation magnitude reverses above subset size 5 (Supplementary Figure 4), making negative local O-information in smaller subsets a special case. As such, a thorough analysis of larger subsets is needed to complete our rough picture of time-varying higher order interactions, which we turn to next.

### D. Optimization for Larger Subsets

The space of possible subsets grows combinatorially larger with subset size. This prohibits exhaustive calculations of all subsets of a given size, as was performed for subsets of size 3 and 4. To address larger subsets we performed an optimization employing a simulated annealing algorithm on every time point (see Methods) to search for subsets with high local synergy or high local redundancy. There is good reason to believe that local O-information values cannot be compared across subset sizes, since the exact value of the local O-information is directly dependent on subset size (see Supplementary Text 1). As such, simulated annealing was run for each subset size independently on all time points. We additionally caution against any interpretation of these results that makes direct comparisons between subset sizes on the basis of O-information magnitude.

Results from the simulated annealing algorithm further indicated that time series of optimally synergistic nodes in smaller subsets (*≤* 15) are often distinct from their counterparts in larger subsets (*>* 15). The percent overlap between subsets of different sizes found at the same time point is shown in Figure 5a, indicating generally lower overlap for smaller subsets. To determine whether nodal overlap was statistically significant, every time point was compared to a null distribution of 1,000,000 samples formed by randomly sampling two subsets of the appropriate sizes and recording the overlap. The fraction of significantly overlapping time points after stringent Bonferroni correction is shown for all subset size pairs in Figure 5b. Little to no significantly overlapping time points are seen for subset size 15 and below. The paucity of significantly overlapping time points between smaller and larger subsets clearly indicate distinct patterns of synergistic interaction on different interactions orders.

**FIG. 5.**
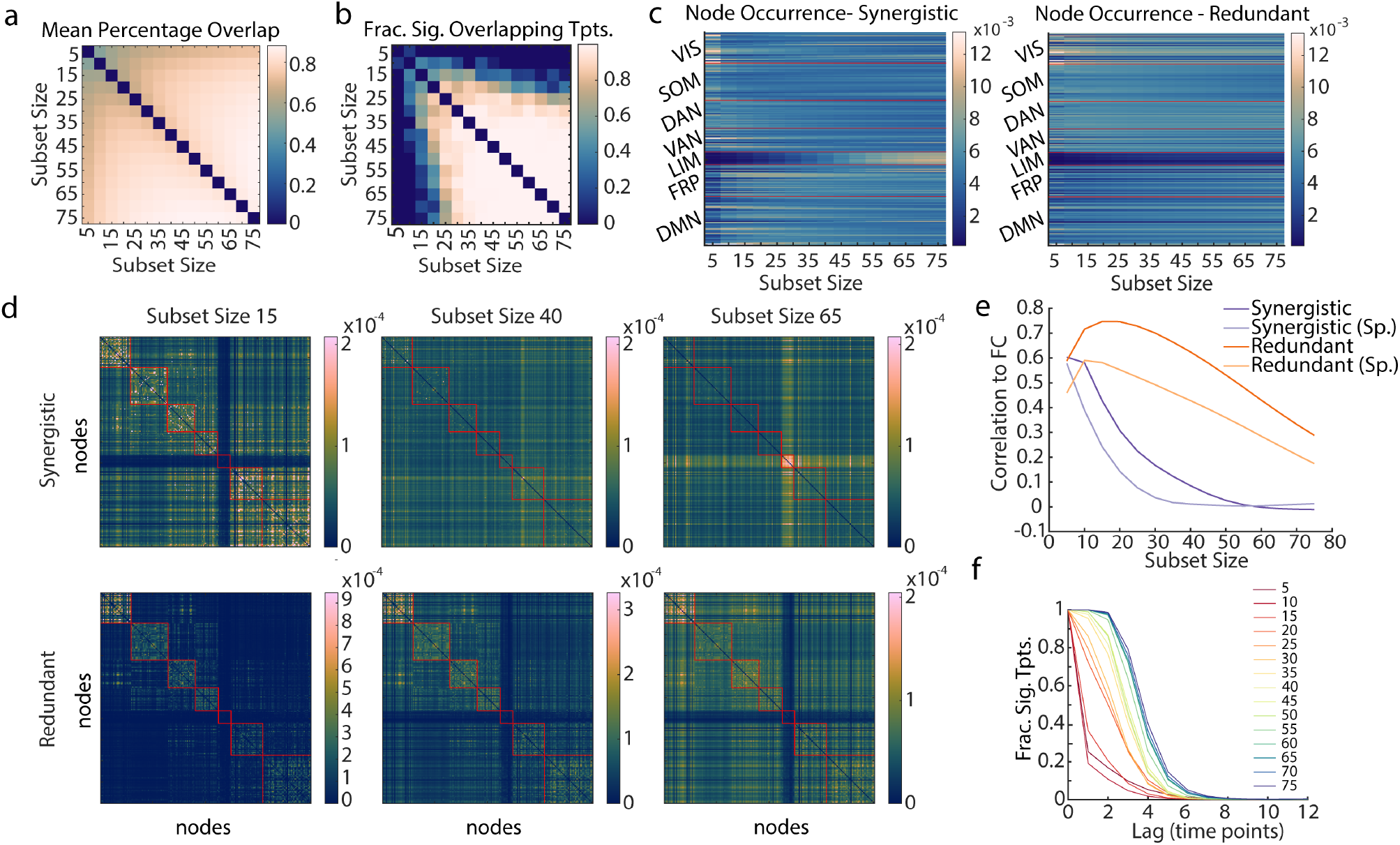
Annealing produces distinct subsets for synergy and redundancy as well as subset size. **a**. The percent overlap of nodes between the same time point optimized at different subset sizes was calculated. Percentages were taken as a fraction of the smaller subset size. Here we show the mean percent overlap between every subset size. **b**. The fraction of time points with significant nodal overlap (*p <* 0.05, Bonferroni corrected, in comparison to a randomly sampled null) between subset sizes, shown for all pairs of subset sizes. **c**. The frequency of node occurrence in synergistic and redundant subsets across all time points, shown for all subset sizes and organized by Yeo system (red lines demarcate systems). **d**. Frequency of nodal co-occurrence in synergistic and redundant subsets for all time points, shown for three sample subset sizes, 15, 40, and 65. Rows and columns are organized by Yeo systems (red lines demarcate systems), in the same order as **c**. See Supplemental figure 5, for all subset sizes. **e**. The Pearson and Spearman correlations of the co-occurrence of nodes in redundant and synergistic subsets to time-averaged functional connectivity across subset size. **f**. The fraction of time points with significant overlap (*p <* 0.05, Bonferroni corrected, in comparison to a randomly sampled null) to time points with lag reported on the x-axis.

However, for smaller subsets (sizes 5-15), the finding that globally redundant subsets exhibit the most synergistic moments tended to hold. In the node occurrences shown in Figure 5c, it can be seen that the Visual and Default Mode Networks are well-represented, as they were in the triads, up to Size 15. The trend changes above this size, and at much larger sizes, nodal participation in synergistic subsets is much most pronounced in the limbic system (Figure 5c). This system is not included at all in smaller synergistic subset sizes. More consistency across subset size is seen in redundancy-optimized subsets. In contrast with synergistic subsets, nodes from the visual system consistently occur, and nodes from the limbic system are almost never chosen. Both of these nodal occurrence patterns were replicated in the MICA dataset (see Supplemental Figure 5)

Plotting the co-occurrence of nodes in annealed subsets (Figure 5d, Supplemental Figure 6 and 7) further supports the idea that only at smaller interaction orders globally redundant subsets have the most synergistic moments. At subset sizes of 15 nodes or less, the co-occurrence matrices from synergy-optimized subsets and redundancy-optimized subsets are equally strongly correlated to the data’s functional connectivity (Figure 5d,e), a structure known to be dominated by globally redundant interactions [10, 12]. This correlation quickly falls off for larger synergistic subsets as the co-occurrence matrices become dominated by the presence of nodes from the limbic system (Figure 5d,e). In redundancy-optimized subsets, the correlation to functional connectivity remains remarkably strong for all sizes, peaking at 0.75 (*p* = 0), and visually the correlation matrices notably echo the functional connectivity matrix (Figure 5d,e). The co-occurrence matrices calculated from optimized MICA subsets showed similar trends in both redundant and synergistic subsets (see Supplemental Figure 5).

**FIG. 6.**
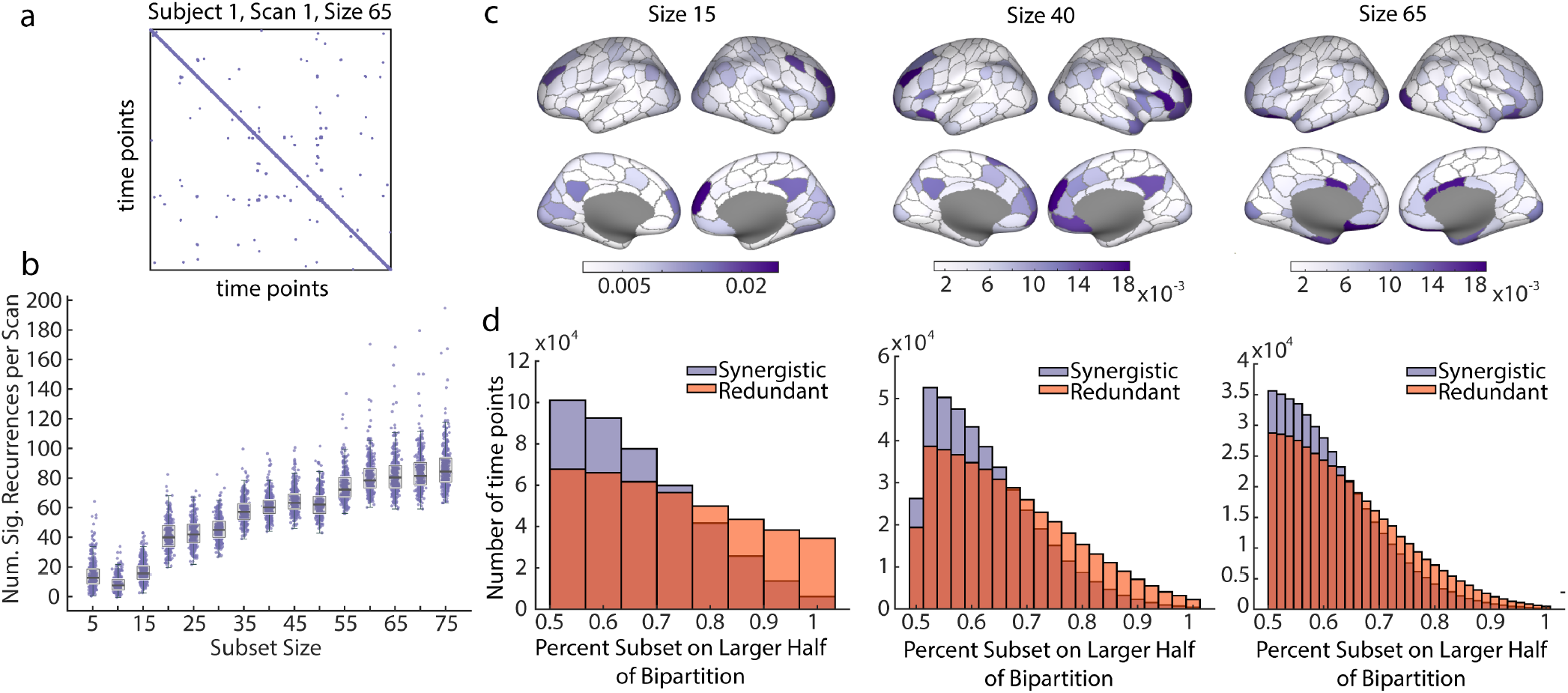
Significant recurrences are found at all subset sizes. Synergy and redundancy dominated subsets have distinct momentary bipartition ratios. **a**. Sample recurrence matrix from subject 1, scan 1 at subset size 65. Purple points represent significant recurrences. **b**. Boxplot showing the number of significant recurrences per scan by subset size. Significant recurrences are present in all subset sizes. **c**. Frequency of nodal participation in significant recurrences, shown for three sample subset sizes, 15, 40, and 65. All subset sizes can be found in Supplemental Figure 6. **d**. Histogram of momentary bipartition ratio (see Results and Methods) for synergistic and redundant subsets, shown for three sample subset sizes, 15, 40, and 65. All subset sizes can be found in Supplemental Figure 7. Redundant and synergistic distributions are significantly different for all subset sizes (Two-way Kolmogorov Smirnov Test, size 15: K-S Stat = 0.1891, size 40: K-S Stat = 0.1452, size 65: K-S stat = 0.1017, all *p <* 0.05, Bonferroni corrected).

We turn to the temporal structure of synergy and redundancy optimized subsets next, beginning with temporal overlap (autocorrelation) of the nodal time series. The fraction of significantly overlapping time points (as compared to a randomly sampled null) is reported at increasing time lags in Figure 5f for all subset sizes. As seen in the triads and tetrads, the subsets involved in synergistic and redundant interactions change relatively smoothly. Larger subsets have greater autocorrelation, but the temporal overlap has completely fallen off by a lag of eight time points for all subset sizes. However a smooth transition between optimized subsets was not seen in the MICA dataset (Supplemental Figure 5). A similar non-replication was seen for the triads and tetrads and may point to a possible sensitivity to scanning parameters.

As in the triads and tetrads, we found significant temporal recurrence of synergistic structures in larger subsets. A significant recurrence was defined as any two time points within the same subset size with statistically significant nodal overlap in comparison to a distribution of 1,000,000 paired samples of random subsets. P-values were Bonferroni corrected to account for all comparisons within a subject, including all subset sizes. In order to avoid conflating the noted autocorrelation of the nodal time series (Figure 5e) with recurrence, only time points at least ten time points apart were considered significant recurrences. Figure 6a shows a sample recurrence matrix for subject 1, scan 1 and subset size 65. Purple points indicate time points with significant recurrence. All subset sizes showed significant recurrence in most scans (Figure 6b).

To understand which nodes tend to recur, we counted the frequency that each node participated in a recurrent time point across the entire dataset. In order to be counted, a node must not only be selected at a time point marked as significantly recurrent, it must also be shared by both the initial and recurrent time point. Frequencies of nodal participation in recurrences are reported on the cortical surface for small (15), medium (40), and large (65) subset sizes in Figure 6c for the HCP data and Supplemental Figure 5 for the MICA data. These plots are meant to be considered analogously to the recurrence plots shown for the triads and tetrads in Figure 3e, and smaller subset sizes replicate many of the regions shown in that plot. However, there were not enough significantly recurrent time points to do a comprehensive clustering analysis, as was performed for the triads and tetrads. Recurrence participation plots for all subset sizes can be found in Supplemental Figure 8.

Finally, a triad or tetrad’s momentary relationship to the bipartition was key for determining whether the subset was synergy or redundancy dominated in the same moment. We found that the same effect was robustly true in optimized subsets of all sizes, small and large alike. To show this, we recorded the larger number of selected nodes with the same z-score sign as a fraction of the total number of selected nodes on every time point (what we call a subset’s “bipartition ratio”). A bipartition ratio of 0.5 indicates that exactly half of the nodes have a positive z-score at that time point while a bipartition ratio of 1 indicates that all nodes have the same sign. The distributions of bipartition ratio for subset sizes 15, 40, and 65 are shown in Figure 6d, and for all subset sizes in Supplementary Figure 9. Results from the triads and tetrads suggest that the fraction of selected nodes on the larger half would tend toward one for momentarily redundant subsets and toward 0.5 for momentarily synergistic subsets. This was exactly the pattern followed for all subset sizes. Synergistic subsets tended to be more evenly split across the bipartition and redundant subsets tended to have entirely the same sign. All synergistic and redundant distributions of the same subset size were significantly different (Two-way Kolmogorov Smirnov test, statistics reported in figure legends, *p <* 0.05, Bonferroni corrected). We replicated this result in the MICA data, but only up to subset size 25 (Supplemental Figure 5). Above that size, the trend reversed, although not strongly.

## IV. DISCUSSION

Here we have provided a novel analysis of how redundancy/synergy dominance evolves through time in resting state BOLD data. This analysis extends and comments on previous studies using the expected value of the O-information on fMRI time series. Focusing on momentary synergy and redundancy dominance reveals that it is redundancy-dominated subsets that experience the most strongly synergy-dominated moments. We have further shown that these synergistic moments often occur when the subset tends to be evenly split across the bipartition created by the signs of the nodal activity, which, for small subsets, occurs at those moments when typically highly coherent systems disintegrate. In addition, we have demonstrated that strongly synergistic and strongly redundant interactions of all sizes have temporal structure. They change smoothly in time and recur at significantly higher rates than expected by chance. Further, we replicated many findings in an independent dataset.

The finding that small (3-20 nodes) redundancy-dominated subsets have the greatest dynamic range in their local O-information values sheds new light on many studies using the expected value of the O-information to identify synergistic interactions. In particular, Varley and Pope et al.[10] used simulated annealing to find subsets with highly negative time-averaged O-information. We expect that the highly synergistic subsets found in Varley and Pope et al. will exhibit muted local O-information dynamics similar to the randomly sampled subsets. While our results did indicate that this relationship may not hold for all subset sizes, the subset sizes explored in Varley and Pope et al. fall within the range where this relationship is quite strong. Using the local O-information as we did in this study will allow more specificity regarding the timing of synergistic and redundant interactions in future studies, and the contrast of our results with those reported in previous work make it clear that temporal specificity is necessary because it determines which interactions are identified.

Increased temporal specificity will also greatly benefit future studies seeking to study synergistic or redundant interactions in cognition or behavior. Temporal specificity will allow elucidation of the exact time-courses of possible higher-order responses to stimuli, just as node-level responses to tasks have been discovered in the past. This is especially relevant to cognitive and behavioral studies given the link between redundancy and and stimulus discrimination [16] and the link between memory task execution and synergy [26]. Further linking synergistic and redundant interactions to cognition would be a fruitful avenue for future research.

In light of the result that redundancy dominated subsets exhibit the most extreme local O-information values, we suggest that it may be more appropriate to think of some subsets of brain regions simply as information-rich, rather than restrictively synergy or redundancy-dominated. We found that for small subsets, and especially in the optimal triads, these information-rich regions correspond to known functional systems, such as the visual system, the somatomotor system and the default mode network. While past work, and indeed most functional connectivity analyses, have focused on the redundant interactions within these systems [12, 17], our results indicate that a thorough exploration of synergistic interactions within and between these systems is necessary. Since it is well-established that many functional systems have strong relationships to cognition, teasing apart the moments in which those systems are interacting synergistically may provide new windows into cognitive processes.

We also showed that often what distinguishes a synergistic moment for a redundancy dominated subset is that subset’s instantaneous relationship to the bipartition. We found this relationship for all subset sizes in the HCP data, and in subset sizes less than 30 in the MICA dataset. Non-replication for larger subsets may be attributable to the much shorter length of the MICA time series. Fewer data points will lead to undersampling of the multivariate joint distribution, and this effect will become more pronounced for larger subset sizes. The robustness of this result in small subset sizes highlights the necessity of synergistic interactions to combine separate streams of information. In this case, we have approximated independent information by the sign of the z-score, which defines two separate momentarily co-fluctuating sets of brain regions. Hence, these two sets represent independent and internally redundant information that is integrated into a synergistic mix. While there are certainly more sophisticated methods for identifying independent sources of information, the bipartition has proved to be a useful heuristic in this case. The relationship of synergistic moments to the bipartition does accord with previous results that in order to be considered synergistic nodes in a subset must be chosen from multiple functional systems [10]. At each moment in time, the most simple division into separate functional systems is into co-fluctuating communities. Subsets that cross these communities will likely be synergistic. The known Yeo functional systems emerge from time-averaging many moments of co-fluctuation [61], and so represent structures that appear on longer time scales. Our results, coupled with those of previous work, may suggest a multi-scale temporal structure of synergistic interactions: short-term synergy occurs within functional systems, and long-term synergistic interactions take place between functional systems.

The relationship to the bipartition is also fortuitous from a computational perspective. It is easily accessible, requiring minimal calculation. Assessing the instantaneous bipartition ratio of a candidate subset before calculating its local O-information time series could improve computation time dramatically and significantly narrow the search space of possible higher-order interactions. Simply, if a researcher wants to study synergistic interactions and samples a subset with a bipartition ratio near one, then it is unlikely to be synergistic and the local O-information need not be calculated. The converse would work for researchers seeking to sample redundant interactions. An important caveat is that because of the non-replication in large MICA subsets, this should only be done for smaller interactions, unless the data has a very large number of time points.

One of the major limitations of this work is that the O-information is a statistical measure, and does not guarantee insight into the underlying higher order mechanisms [62]. Certainly, our results do indicate the possible presence of higher-order mechanisms, but further studies should attempt to bridge this gap. Both dynamical modeling approaches [63] and more thorough examination of anatomical structure within and between the key regions [32] will be of use in this effort. This limitation is further exacerbated by the fact that we have applied the local O-information to BOLD data. Since the BOLD signal is a hemodynamic response to underlying neuronal activity and not the activity itself, we are further limited in our ability to make claims about neuronal mechanisms. Additionally, the BOLD response is slow and temporally offset due to the slow time course of the hemodynamic response function [64]. However, our results may provide some suggestion of the kinds of temporal structure other researchers may expect to see when applying the local O-information to their own work. An opportunity for future exploration would be to apply the methods employed here to data from smaller scale neuronal population recordings. Some studies have already made use of synergy at this scale [22, 23, 65], and the local O-information represents an opportunity to continue these lines of research and expand them to larger interactions.

However, as some of the results in the current work indicate, using the O-information to study very large interactions must be done with caution. The largest interaction we considered in this paper was at the level of the whole brain. Synergistic interactions were exceedingly rare and produced framewise functional connectivity matrices with very little structure. Relatedly, large subset sizes (*≥* 55) heavily incorporated the limbic system, a system known to have poor signal to noise ratio in BOLD data. This may indicate that the local O-information is highly sensitive to noise, a possibility which has been previously reported in the literature for the time-averaged form of the O-information [66]. We suggest that the local O-information’s sensitivity to noise may be exaggerated when it is used on subsets that are larger than the true scale of interaction for the system. Two lines of reasoning support this suggestion. First, the inclusion of highly noisy nodes in our results only happened at very large subset sizes. Second, previous results [10, 12], as well as the results relating subsets to the bipartition in this work, indicate that subsets tend to be synergistic when they incorporate multiple sources of independent information. At a small subset size this can be accomplished by selecting nodes that are uncorrelated. At larger subset sizes this becomes harder as there is a limited number of independent information streams in the data, and so some correlated nodes must be selected in order to meet the required size. When this happens, incorporating noise may also be a way of introducing independent information to the subset. Low signal to noise ratio regions will often vary erratically in relation to the rest of the brain, providing reliably independent signals against the backdrop of otherwise largely correlated functional systems. Incorporating these noisy nodes may be an easy way for our optimization algorithm to keep the O-information low when the subset size is large enough to force the inclusion of many correlated nodes. Of course, the best solution to this problem is to use data with as little noise as possible. But when this is not possible, it will be paramount that the interpretation of a negative value of the local O-information be done with care and with an eye to the possibility of conflation of noise with synergy.

Finally, our most striking and promising result, was the significant temporal structure found across all subset sizes. We found remarkable recurrence in the optimal triads and recurrence was also a prominent feature in the optimized larger subsets. While the subsets found by optimization are certainly limited by non-exhaustiveness, this fact also makes the significant recurrences even more salient. The presence of these features in our non-exhaustively optimized data indicate that they must be quite a strong feature of the data in reality. Given much more time and computational power we can be fairly certain that many more significant recurrences would be found. The presence of recurrence indicates that the interactions our analysis identified are not random, but represent states of activity that the cortex repeatedly returns to. This suggests that these interactions may play a significant role in cognition, or may at least serve in the identification of brain states by researchers seeking to study cognition.

As a first application of the local O-information to BOLD fMRI data, we have shown that it is effective in identifying highly recurrent interactions with specific internodal relationships. Both of these traits make it a promising measure for future application to studies of on-going cognition. We hope that the study of time-resolved synergistic interactions may provide a new look at the neural underpinnings of many cognitive processes.

## V. ACKNOWLEDGEMENTS

The authors would like to thank Maria Grazia Puxeddu for many helpful discussions and feedback during the process of preparing this manuscript. M.P. is supported by the National Science Foundation Graduate Research Fellowship under Grant No. 2240777. Any opinion, findings, and conclusions or recommendations expressed in this material are those of the authors and do not necessarily reflect the views of the National Science Foundation. This research was supported in part by Lilly Endowment, Inc., through its support for the Indiana University Pervasive Technology Institute which provided the computer infrastructure required for completion of the project. The funders had no role in study design, data collection and analysis, decision to publish, or preparation of the manuscript. Data were provided, in part, by the Human Connectome Project, WU-Minn Con sortium (principal investigators: D. Van Essen and K. Ugurbil; 1U54MH091657), funded by the 16 NIH Institutes and centers that support the NIH Blueprint for Neuroscience Research and by the McDonnell Center for Systems Neuroscience at Washington University. M.P. would additionally like to thank her husband for his unending support.

