## Supplemental Figures and Text for "Time-varying synergy/redundancy dominance in the human cerebral cortex"

### Supplemental Information

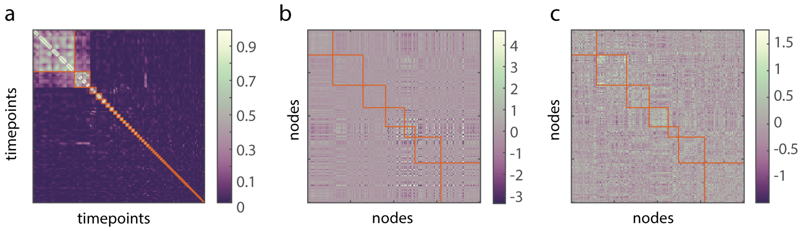


FIG. 1. **Communities of whole brain synergistic moments are noisy. a.** A matrix of correlations between the co-activation matrices of whole brain synergistic time points, organized by the communities found by multi-resolution consensus clustering. Most time points did not fit well within a community. **b.** The centroid of the co-activations of the largest community of whole brain synergistic time points. Yeo systems are marked with orange boxes. **c.** The centroid of the co-activations of the second-largest community of whole brain synergistic time points. Yeo systems are marked with orange boxes. Neither co-activation matrix shows much structure.

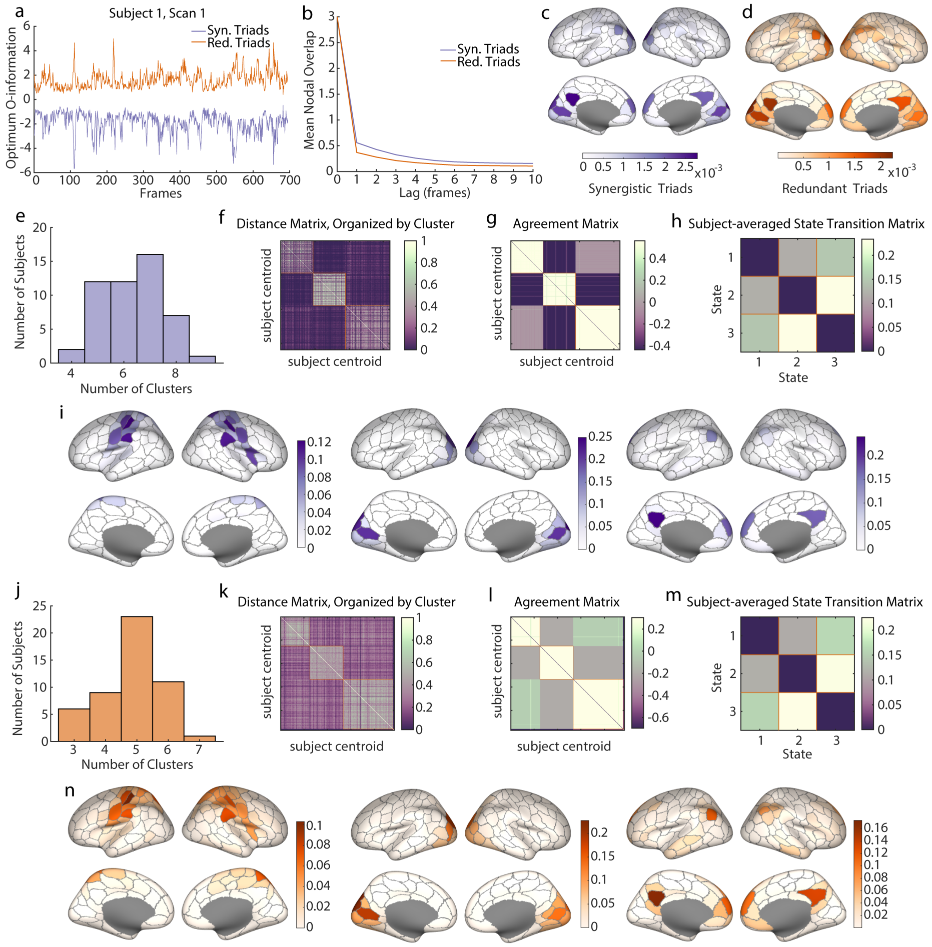


FIG. 2. **Optimally redundant and synergistic triads in the MICA dataset replicate nodal participation and recurrence patterns. a.** A sample time series of the maximum and minimum local O-information found in the MICA dataset at each time point, which may act as upper and lower bounds of the local O-information. **b.** The average number of nodes shared between time points at varying frame lags. This serves as an autocorrelation for the nodal time series. **c.-d.** The frequency of participation in synergistic and redundant triads plotted on the surface of the cortex. This replicates the same pattern of participation found in the HCP data. **e., j.** A histogram of the number of recurrence clusters found in each subject during subject-level multi-resolution consensus clustering in synergistic and redundant triads, respectively. **f., k.** Matrix of distance between subject centroids found in the subject-level multi-resolution consensus clustering. The matrix is ordered according to the resulting group level clusters. **g., l.** Agreement matrix reporting the frequency with which subject centroids were assigned to the same cluster during multi-resolution consensus clustering. Also ordered according to the resulting group level clusters. **h., m.** Group level clustering resulted in three main clusters (states) for both synergistic and redundant triads (redundant triads had an additional cluster composed of three time points that is not reported here). The unique assignment of each time point to a state allows the transitions between each state to be recorded within a single scan. The frequency of transition is then averaged across subjects and reported in state transition matrices. **i., n.** Frequency of nodal participation in synergistic and redundant recurrent structures, respectively. These nodes are more frequently selected together as participating in most synergistic or redundant triads.

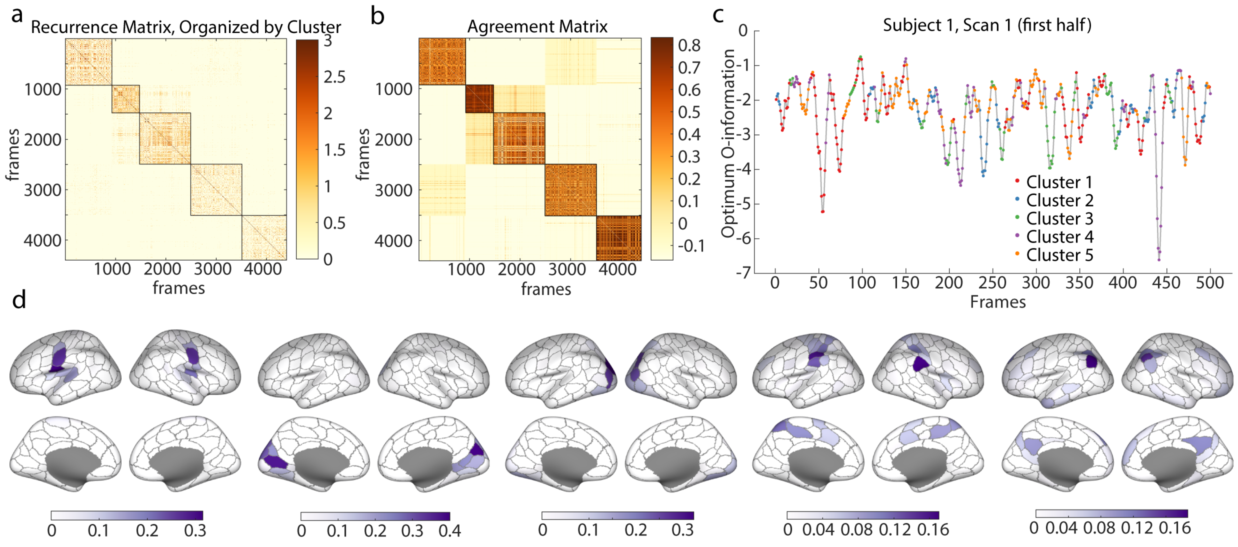


FIG. 3. **Single-subject recurrent patterns.** **a.** The recurrence matrix of all four scans of the first subject. Values in the matrix are jaccard similarity scores between subsets chosen on each frame. Rows and columns have been organized according to the five clusters found with multi-resolution consensus clustering (MRCC). **b.** Agreement matrix recording community co-assignment of frames during MRCC. **c.** A sample time series of optimal local O-information values from subject 1's first scan, where points have been colored by the five recurrence structures, showing good temporal contiguity. **d.** Nodal participation frequencies in each of the five recurrence clusters. Frequencies are calculated as the number of times a node appears in subsets chosen at the time points belonging to a given cluster divided by the number of time points in the same cluster.

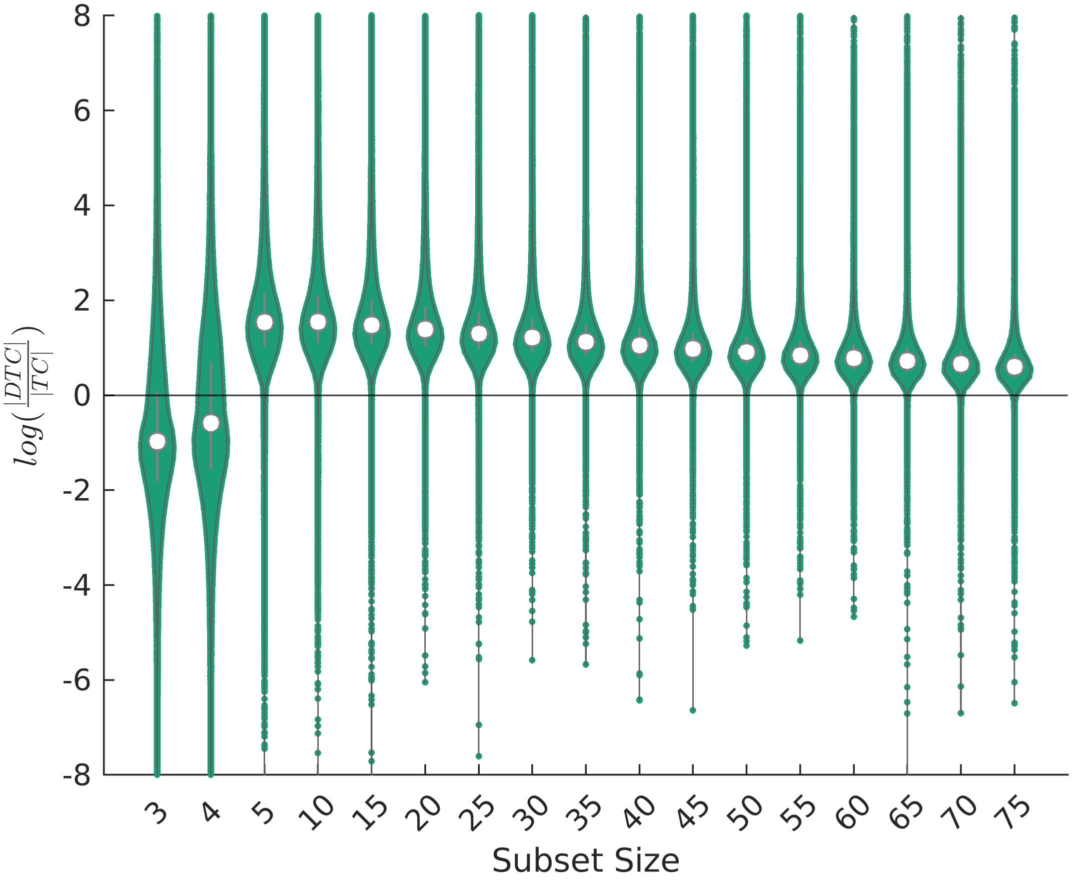


FIG. 4. **Negative local O-information is differentially driven by TC and DTC at different subset sizes.** Because localized TC and DTC can take on negative values, local O-information can be negative either because of a strong positive value in local DTC or a strong negative value of local TC. Here we show violin plots of the log of the ratio of the magnitude of the local DTC to the magnitude of local TC at different subset sizes. Where the log of the ratio is below zero, the local O-information is negative because of strongly negative local TC values. Where the log of the ratio is above zero, the local O-information is negative because of strongly positive local DTC values. The plot shows a clear shift from high magnitude TC to high magnitude DTC between subsets size 4 and 5, suggesting that small subsets may have particular properties not readily generalizable to large subsets. For an in-depth explanation of negative local TC in BOLD fMRI data, see Supplemental Text 1 below.

##### Supplemental Text 1

There are several conditions under which the local O-information can be negative. Since the local O-information is given by $\omega(\mathbf{x}) = tc(\mathbf{x}) - dtc(\mathbf{x})$, negativity can occur when $tc(\mathbf{x}) < 0$ and $|tc(\mathbf{x})| > |dtc(\mathbf{x})|$, or when $dtc(\mathbf{x}) > 0$ and $|tc(\mathbf{x})| < |dtc(\mathbf{x})|$.

Negative local total correlation is the most frequently occurring condition in optimally synergistic triads and tetrads. These moments occur when a subset that is usually strongly coherent (such as a Yeo system) becomes incoherent. Here, using the mathematics of the local total correlation, and what we know about the particular structure of BOLD fMRI data, we demonstrate why that must be the case.

The local total correlation is given by:

$$tc(\mathbf{x}) = \sum_i h(x_i) - h(\mathbf{x}) \tag{1}$$

where $h(x_i)$ is the local entropy for a given state of the variable $x_i$, or the Shannon surprisal $(-log(p(x_i)))$ of that state. We are interested in the case that the local total correlation is negative, or

$$0 > \sum_i h(x_i) - h(\mathbf{x}). \tag{2}$$

Rearranging this equation and writing the surprisal in terms of the log probabilities of the state, we can write:

$$log(\frac{1}{p(\mathbf{x}}) > \sum_i log(\frac{1}{p(x_i)}). \tag{3}$$

Then, exponentiating both sides, this becomes:

$$p(\mathbf{x}) < \Pi_i p(x_i). \tag{4}$$

In other words, the local total correlation is negative when the joint probability of a state is less than the probability of seeing the same state if each unit were acting independently.

If a system spends most of the time coherent (such as a Yeo system), this condition is particularly easy to satisfy. To see why this is the case, we will first consider a Gaussian system that has been binarized at its mean and then consider the same case but in the full continuous space. The principle does not change when we move into the continuous space, but it is easier to see in the binary. For a binarized Gaussian system, each variable $x_i$ spends equal amounts of time above and below its own mean, meaning that in the limit of infinite time, the probabilty of observing the variable in either state is $p(x_i) = 0.5$. Thus we can say that for any Gaussian system binarized about its mean:

$$\Pi_i p(x_i) = 0.5^i \tag{5}$$

Inserting this into equation 4, we see that the local total correlation will be negative when

$$p(\mathbf{x}) < 0.5^i. \tag{6}$$

In other words, the local total correlation will be negative whenever a subset enters a state that is relatively rare for that subset. For a subset of size 3, the joint state must occur less than one-eighth of the time in order to satisfy this condition. Of course, this is dependent on subset size. The larger the subset, the more unlikely a joint state would have to be to result in a negative local total correlation.

For a system to satisfy this requirement some of the time, the distribution of joint states must be non-uniform such that some joint states are visited far more frequently than others. In the case of the brain, it is known that many sets of brain regions corresponding to traditional resting state networks spend a large portion of their time integrated, creating a highly non-uniform distribution of joint states, in which the integrated joint states are attributed a much larger probability mass than their distintegrated counterparts. It is easy to see that the greater the difference between the probability of the joint state and the probability of the independent case, the more negative the local total correlation will become. Said another way, the more the mass of the distribution of joint states is concentrated on only a few joint states, the more negative the local total correlation will become when rare states appear.

Returning to the continuous condition, because our data is z-scored we can assume the single variables follow a standard normal distribution with $\mu = 0$ and $\sigma = 1$. The most likely value for a random variable following this distribution to assume is $\mu$, which has probability density $\frac{1}{\sqrt{2\pi}}$. We can substitute this as an upper bound for the probability density of a single state of the random variable $x_i$ in the right hand side of equation 4, so that

$$p(\mathbf{x}) < \Pi_i p(x_i) < (\frac{1}{\sqrt{2\pi}})^i \tag{7}$$

Again, this leads us to the result that the more rare the joint state of the variables, the more negative the local total correlation will be. In the case of triads, $p(\mathbf{x}) <$ 0.0635 for the total correlation to be negative.

This also helps explain why a negative log ratio of the magnitude of local DTC to local

TC at time points with negative local O-information becomes unlikely at larger subset size (see Figure4). The joint state must be increasingly less likely as subset size increases in order to have negative local TC.

In general, it is also worth pointing out that equations for both local TC and local DTC depend directly on the subset size, in the manner of equation 4. This renders direct comparison of local O-information values across subset sizes unwise, if not impossible.

**A Note about interpretation**: The fact that the local total correlation becomes strongly negative at rare, unexpected moments may seem like it defeats the notion of synergy. It is worth remembering that the local entropy, or Shannon surprisal, can also be considered the "information content" of a particular state. That this is greater for the joint state than the individual variables acting independently could be considered to match the notion of synergy. Quite literally, it is when "the information content of the joint is greater than the sum of its parts."

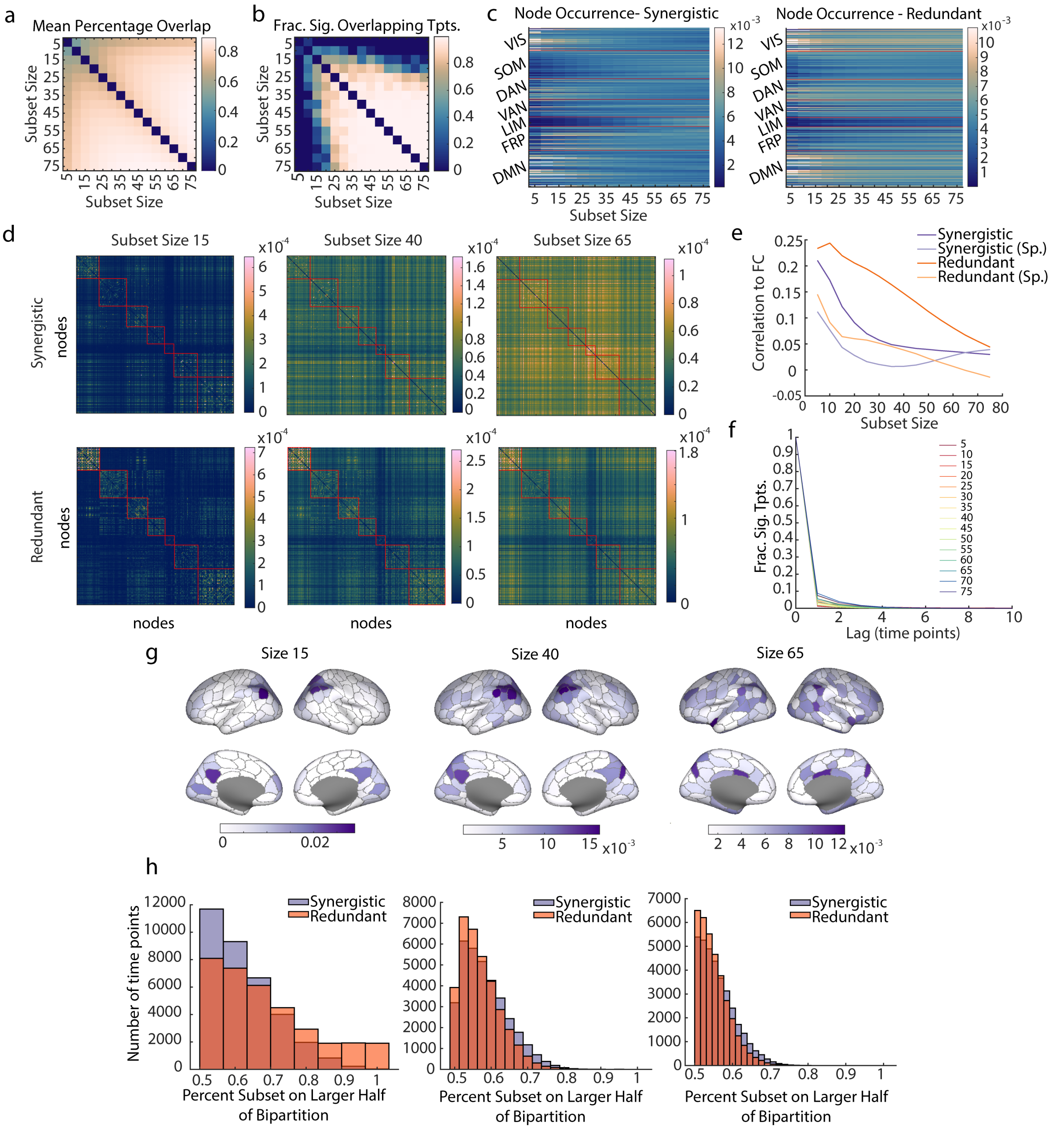


FIG. 5. **Optimized subsets from the MICA data exhibit similar occurrence, co-occurrence, and recurrence patterns to the HCP dataset.** **a.** The percent overlap of nodes between the same time point optimized at different subset sizes was calculated for the MICA data set. Percentages were taken as a fraction of the smaller subset size. Here we show the mean percent overlap between every subset size. **b.** The fraction of time points with significant nodal overlap ($p < 0.05$, Bonferroni corrected, in comparison to a randomly sampled null) between subset sizes, shown for all pairs of subset sizes. **c.** Node occurrences in subsets optimized for synergy and redundancy in the MICA data set. **d.** Co-occurrence matrices for nodes optimized for synergy and redundancy reported for sizes 15, 40, and 65 from the MICA data set. **e.** The correlation of the co-occurrence matrices to the data's time-averaged functional connectivity matrix across subset sizes. **f.** The fraction of time points with significant overlap ($p < 0.05$, Bonferroni corrected, in comparison to a randomly sampled null) to time points with lag reported on the x-axis. **g.** Frequency of node occurrence among recurrent time points in the MICA data set for three subset sizes: 15, 40, and 65. **h.** Histograms of the instantaneous bipartition ratio for synergy-optimized and redundancy-optimized subsets. All differences between distributions are significant (Kolmogorov-

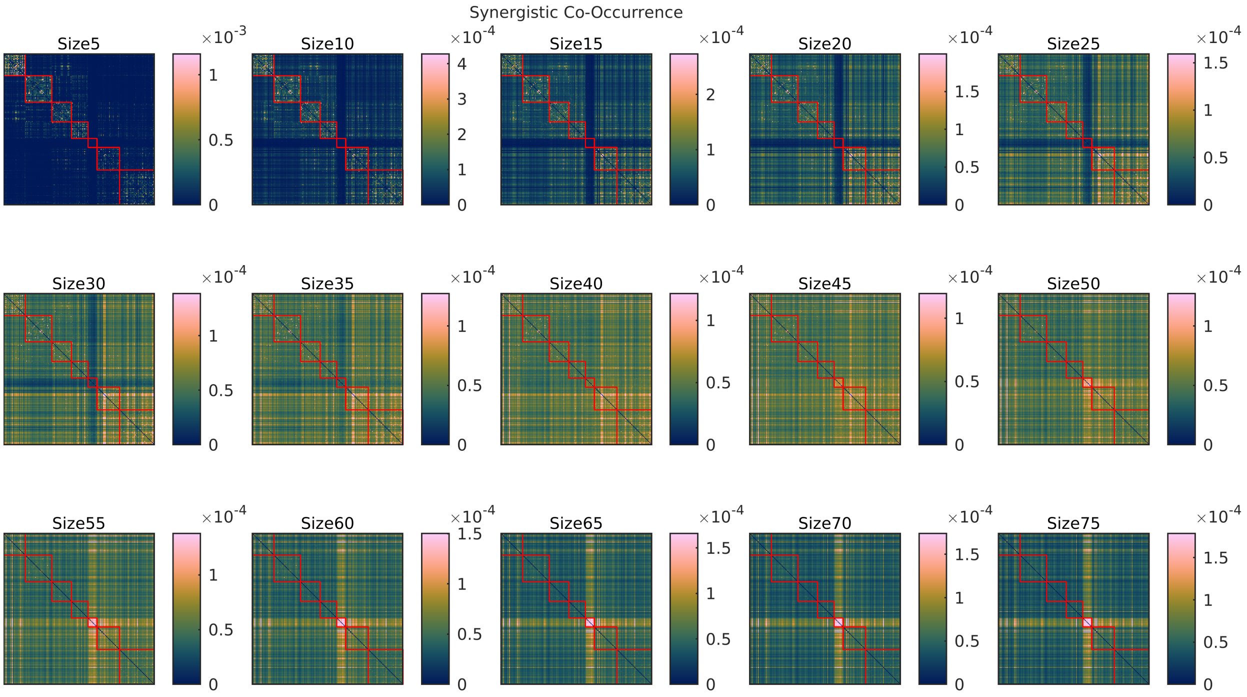


FIG. 6. **Nodal Co-occurrence in synergistic subsets varies with subset size.** Matrices with frequencies of nodal co-occurrence in annealed synergistic subsets are shown for all subset sizes.

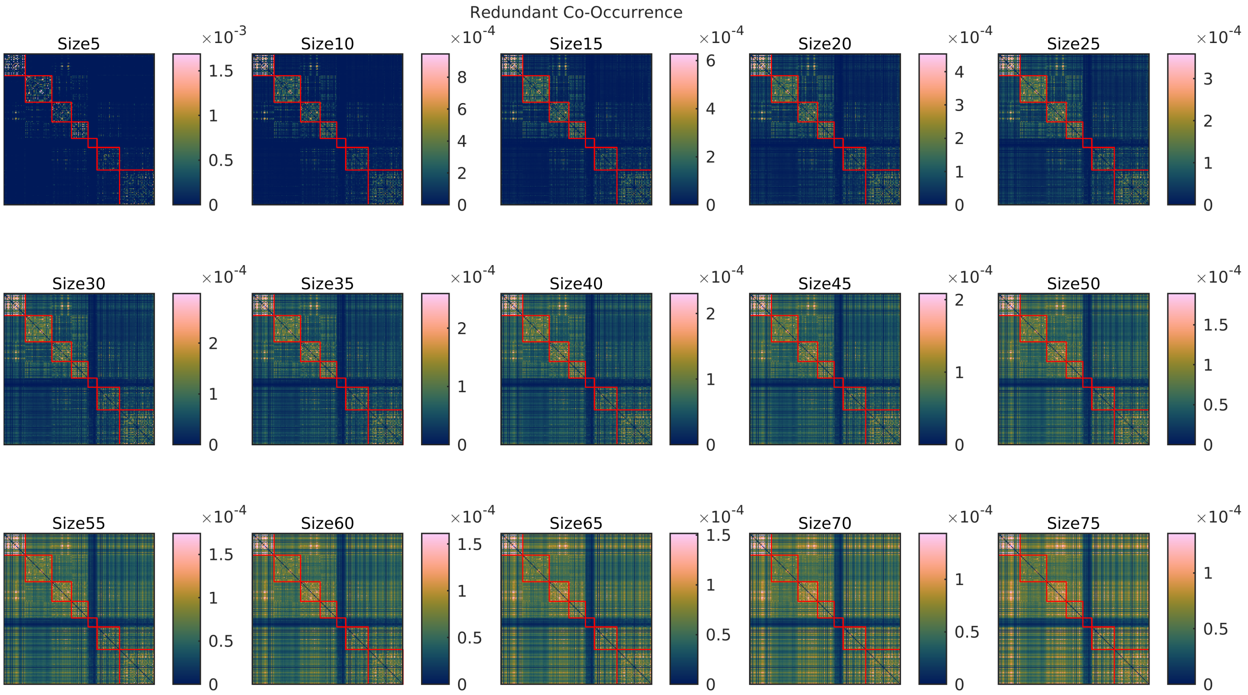


FIG. 7. **Nodal Co-occurrence in redundant subsets varies with subset size.** Matrices with frequencies of nodal co-occurrence in annealed redundant subsets are shown for all subset sizes.

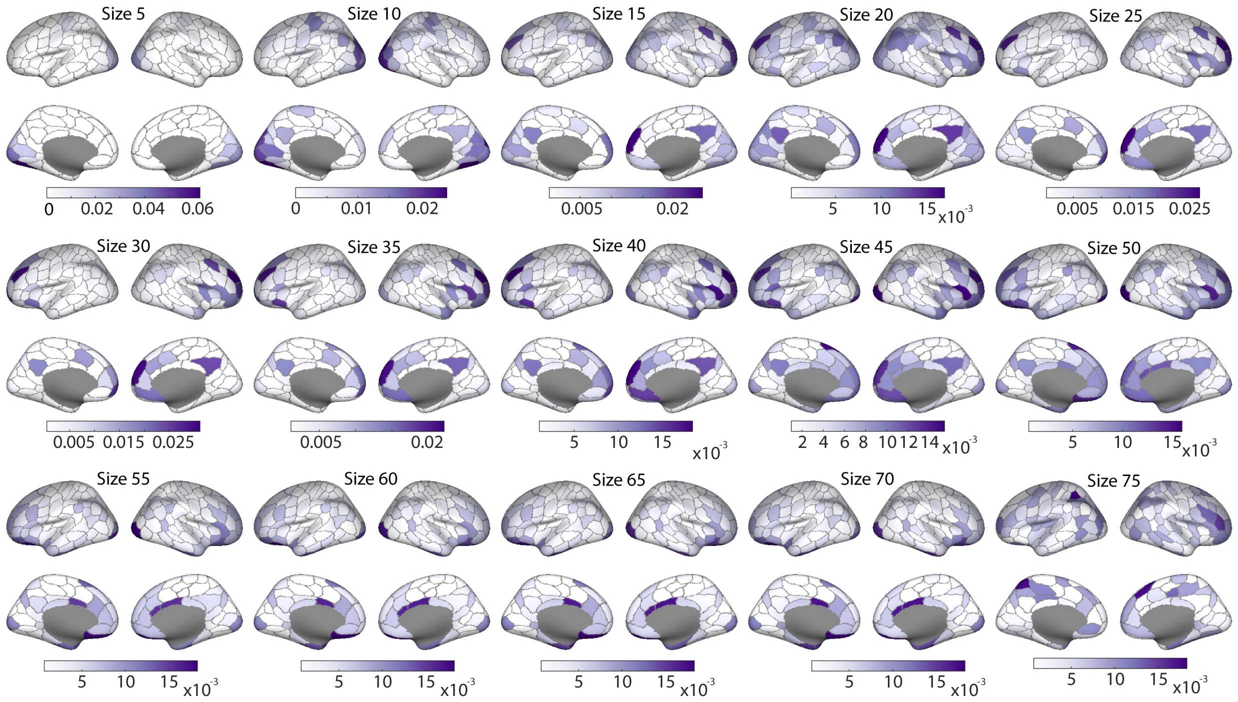


FIG. 8. **Nodal participation in significant recurrences is highly variable across subset size.** Nodes that significantly recur in time are colored on the cortical surface. Intensity of color indicates the frequency of recurrence. Across subset sizes, highly recurrent nodes vary dramatically.

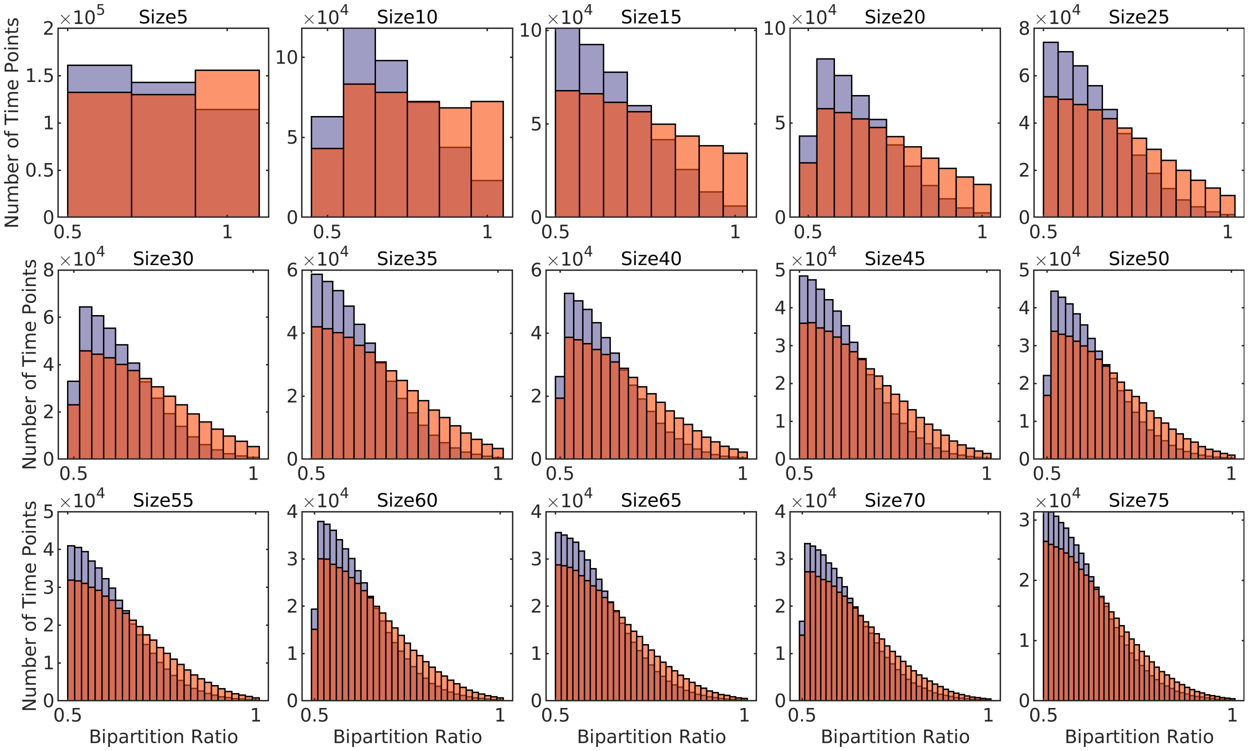


FIG. 9. **The relationship to the bipartition ratio remains the same, across subset sizes.** Histograms of the bipartition ratio for redundant and synergistic subsets are shown for all subset sizes studied. Regardless of subset size, synergistic subsets tend toward a bipartition ratio of 0.5 and redundant subsets tend toward a bipartition ratio of 1. All pairs of distributions are significantly different from each other, Kolmogorov-Smirnov two-sample test. size 5: K-S Stat = 0.0994, size 10: K-S Stat = 0.1781, size 15: K-S Stat = 0.1891, size 20: K-S Stat = 0.1838, size 25: K-S Stat = 0.1760, size 30: K-S Stat = 0.1648, size 35: K-S Stat = 0.1542, size 40: K-S Stat = 0.1452, size 45: K-S Stat = 0.1360, size 50: K-S Stat = 0.1265, size 55: K-S Stat = 0.1191, size 60: K-S Stat = 0.1103, size 65: K-S stat = 0.1017, size 70: K-S Stat = 0.0936, size 75: K-S Stat = 0.0851, all $p < 0.05$
